# Distinct Functional cerebral Hypersensitivity networks during incisional and inflammatory pain in rats

**DOI:** 10.1101/2023.01.09.523228

**Authors:** S. Kreitz, B. Pradier, D. Segelcke, S. Amirmohseni, A. Hess, C. Faber, E. Pogatzki-Zahn

## Abstract

Although the pathophysiology of pain has been investigated tremendously, there are still many open questions, especially with regard to specific pain entities and their pain-related symptoms. To increase the translational impact of (preclinical) animal pain neuroimaging studies, the use of disease-specific pain models, as well as relevant stimulus modalities, are critical. Yet, the challenges of identifying neuroimaging signatures at a pain entity- and modality-specific level are manifold. Therefore, we developed a comprehensive framework for brain network analysis in disease-specific pain models combining functional MRI with graph-theory and data classification by linear discriminant analysis. This enabled us to expand our knowledge of stimulus (mechanical vs. electrical) modality processing under incisional (INC) and pathogen-induced inflammatory (CFA) pain entities compared to acute pain conditions. In short, graph-theoretical analyses revealed distinct Network Signatures of Pain Hypersensitivity (NSPH) for INC and CFA, resulting in impaired discrimination of stimulus modalities in both pain models compared to control conditions (CTR). Such specific neuroimaging signatures are an important step toward identifying novel pain biomarkers for certain diseases and relevant outcomes to evaluate target engagement of novel therapeutic options, which ultimately can be translated to the clinic.

## Introduction

Pain is defined as an unpleasant sensory and emotional experience associated with, or resembling that associated with, actual or potential tissue damage (***Raja et al., 2020***). Consequently, pain is a multidimensional process caused by tissue damage, pathogen-induced inflammation, among others, which adjusts behavior to restore and maintain physical integrity and homeostasis. If not resolved, pain can most likely become chronic, leading to additional devastating symptoms (***Smith et al., 2019***). The mechanisms behind pain and pain-related symptoms, in particular its chronification process, are complex and involve multilevel modulations and adaptations. Ultimately, in the central nervous system (CNS), all peripheral and spinal signals are integrated to generate corresponding output signals, of which the underlying mechanisms are accepted to be essential for pain chronification. These processes are, however, complex and not yet well elucidated, especially due to the wide range of diseases leading to chronic pain, with in part different signs and symptoms.

Translational approaches using human (***Burgmer et al., 2012; Maihöfner et al., 2005; Maihöfner and Handwerker, 2005***) and rodent experimental pain models (***Abaei et al., 2016; Amirmohseni et al., 2016***) are necessary to acquire a deeper understanding of the processing relevant for different pain-related symptoms and diseases (***Hansson, 2021***). As shown by using disease-specific pain models such as surgical injury or inflammation, the peripheral (PNS) and the CNS are sensitized in a disease-specific way, resulting in non-evoked and evoked exaggerated pain-related responses resembling symptoms of patients with pain (***Mouraux et al., 2021; Pogatzki-Zahn et al., 2018***). Of these symptoms, mechanical hyperalgesia is prominent and clinically very important, not only for the acute phase but also for chronification of, for example, postsurgical pain, which is a very relevant chronic pain state (***Rosenberger and Pogatzki-Zahn, 2022; Schug et al., 2019***). In patients, mechanical hyperalgesia differentiates postsurgical from neuropathic pain; for the latter, non-evoked pain as well as mechanical allodynia is more prominent (***Mouraux et al., 2021***).

To increase the translational impact of pain neuroimaging studies, the use of disease-specific pain models and relevant stimulus modalities are critical (***Mouraux et al., 2021; Tracey, 2021***). However, despite its frequent use in animal behavioral studies, mechanical stimulation is technically not readily applicable to imaging studies; instead, electrical impulses are most frequently used as a proxy for peripheral sensory stimulation. We recently developed a device evoking calibrated, reproducible mechanical stimuli to rat hind paws in neuroimaging studies to overcome this technical barrier (***Just et al., 2020***). Herewith, we previously studied blood-oxygen-level-dependent (BOLD) fMRI activation of pain-related brain structures caused by mechanical vs. electrical stimulation in disease-specific pain models for incisional and pathogen-induced inflammation pain (***Amirmohseni et al., 2016***). We found that BOLD responses in brain structures differed between pain entities, although the mechanical hypersensitivity assessed by behavioral readouts was similar. However, deriving specific BOLD signatures for pain entities or modalities solely based on BOLD responses of single brain structures was not possible. Even innocuous sensory inputs show large overlaps with noxious stimuli in BOLD-activated brain structures, calling any direct pain specificity *per se* into question (***Amirmohseni et al., 2016; Iannetti and Mouraux, 2010; Legrain et al., 2011; Mouraux et al., 2011***). These findings inevitably lead to the question of whether pain entities and modality-specific neuroimaging signatures can be defined with more advanced analytics (***Wager et al., 2013***).

During the last decade, the idea has risen that specific processing mechanisms rely on functional interactions within complex brain networks (***Bullmore and Sporns, 2009; Sporns, 2011***). In principle, these networks reflect the information flow efficiency between their components and are characterized by topological properties influencing the integration and/or segregation of this information flow (***Sporns, 2013a; Wig, 2017***). Often functional brain networks are assessed in resting-state fMRI, which has been a powerful tool to differentiate between physiological and pathological brain states, including chronic pain (***Baliki et al., 2014; Bilbao et al., 2018; Komaki et al., 2016***) and acute craniofacial pain (***Becerra et al., 2017; Nasseef et al., 2021; Spisák et al., 2017***) in rodents. However, studies of functional brain networks during stimulation states are less frequently used (***Griessner et al., 2018; Neely et al., 2010; Schnorr et al., 2014***). Moreover, we have previously shown such transient effects during acute noxious heat stimulus-evoked BOLD responses in a disease-specific model of arthritis in mice (***Hess et al., 2011***). Graph-theoretical brain networks showed a tight clustering of thalamic and periaqueductal gray (PAG) structures compared to control, which rapidly dissolved after TNF-α antagonist treatment. These findings could be directly translated to human rheumatoid arthritis patients (***Hess et al., 2011; Rech et al., 2013***). Similarly, human imaging studies identified altered functional connectivity (FC) between brain structures restored by pregabalin and let to pain relief in these patients (***Harris et al., 2013***). As another example, an imaging study in healthy human participants recently reported extensive reorganization of brain networks during acute noxious (versus innocuous) heat stimulation, including altered connectivity patterns and distributions of whole-brain hub regions (***Zheng et al., 2019***). Therefore, graph-theoretical network- and connectivity analyzes seem highly relevant for understanding brain processes, even under disease conditions, and for identifying pain entity- and modality-specific brain networks (***Kucyi and Davis, 2017***). But, as a caveat, advanced analyses lead to higher-dimensional data spaces, which are subject to the curse of dimensionality, which requires more sophisticated data science.

In this study, we established an analytical framework as a prerequisite for integrating stimulus-driven BOLD fMRI, graph-theoretical network analyzes, and data classification by linear discriminant analysis. We applied this workflow to pain disease-specific models - incisional and pathogen-induced inflammatory pain - using low- and high-intensity mechanical and electrical stimulation. The network signatures extracted were specifically validated against those from sham-treated animals, reflecting the corresponding acute pain response. This highly comprehensive analysis framework of whole-brain networks enabled us to carve out pain entity- and modality-specific brain network signatures, breaking down specific hypersensitivity signatures at a previously unknown level of mechanistic insight for neuroimaging data.

## Results

The rationale of our approach was the following: we analyzed the brain structure-specific activation probability after peripheral stimulation and calculated the brain-wide functional connectivity. Using graph-theoretical methods, we obtained a detailed view of pain entities and modalities, which enabled us to define brain network signatures of pain hypersensitivity (NSPH). For this purpose, first the individual BOLD MRI data were registered with a digital rat brain atlas, and second, the time courses of significantly activated BOLD voxels in all regions of interest (ROI) evoked by different stimulation modalities (electrical (ES) or mechanical (MS)) at two different intensities (low or high) were extracted (Figure 1a). The pair-wise Pearson’s correlation coefficients of these average time courses per ROI’s represented the functional connectivity between them and build the pain entity-, modality- and intensity-specific networks (Figure 1b). Of note and in agreement with our previous report (***Segelcke et al., 2021b***), both pain models – incisional (INC) and infection-induced inflammatory (CFA) pain – showed similar mechanical hypersensitivity in behavioral tests (Supplementary Figure 1). The networks were assessed by graph-theoretical analysis and all response parameters were subjected to LDA (Figure 1c).

**Figure 1.**
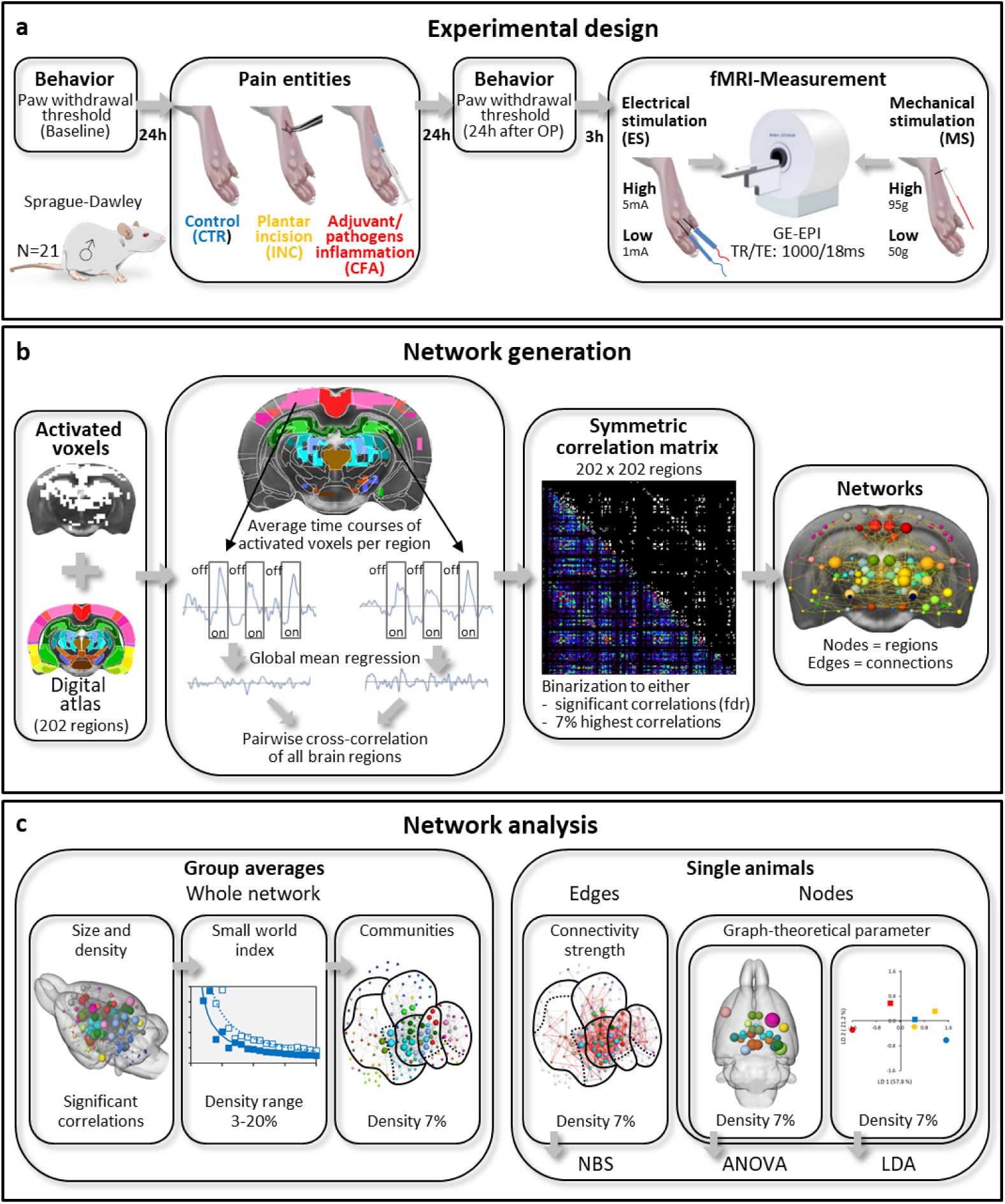
Study design and analysis strategy. **a)** Determination of mechanical threshold was performed prior to (baseline), 24 after pain model induction (plantar incision (INC), adjuvant/pathogen-induced inflammation induced by plantar injection of Complete Freund’s Adjuvants (CFA)), or, as control, brief anesthesia (Isoflurane) only (CTR) in male Sprague-Dawley rats (N=21). fMRI measurements were performed using two different stimulation modalities (ES and MS) with low (1mA / 50g) and high (5mA / 95g) intensity, respectively. **b)** Activated voxels, derived from standard general linear model (GLM) analysis, were assigned to specific brain structures by registration to a digital 3D rat brain atlas. The average time courses of activated voxels per brain structure were extracted, and after regression of their global mean, the residuals were subjected to pair-wise cross-correlation of all activated brain structures. The resulting symmetric correlation map was binarized to all significantly correlating connections (fdr, q=0.05: network-size comparison) or the 7% strongest connections (topology comparison) and subsequently translated into networks with brain structures as nodes and their respective connections as edges. Edge weight, i.e. the correlation coefficient (Pearson’s r), indicated connectivity strength. **c)** Global network characteristics were determined using average networks per group. Statistical analysis of specific connections and nodes was performed using single animal networks. NBS: network-based statistics, ANOVA: analysis of variance, LDA: linear discriminant analysis.

### High stimulation intensity results in bilateral pain entity- and modality-specific, local brain activation

From the previously published statistical parametric map data (***Amirmohseni et al., 2016***) we generated activation probability maps, i.e., the group proportion of significantly activated voxels per brain structure, to indicate stimulus-specific brain activation probability per group. Because of the binary classification of activated versus non-activated voxels, the activation probability is less dependent on the temporal signal-to-noise ratio (tSNR) than the GLM derived t-statistics. Voxel-wise activation probability increased with high-intensity stimulation for all pain entities and both stimulation modalities. The total activated brain volume with high probability was higher for both pain models than CTR (Figure 2a), but increased intensity-dependent only with INC and CTR, not in the CFA group.

**Figure 2:**
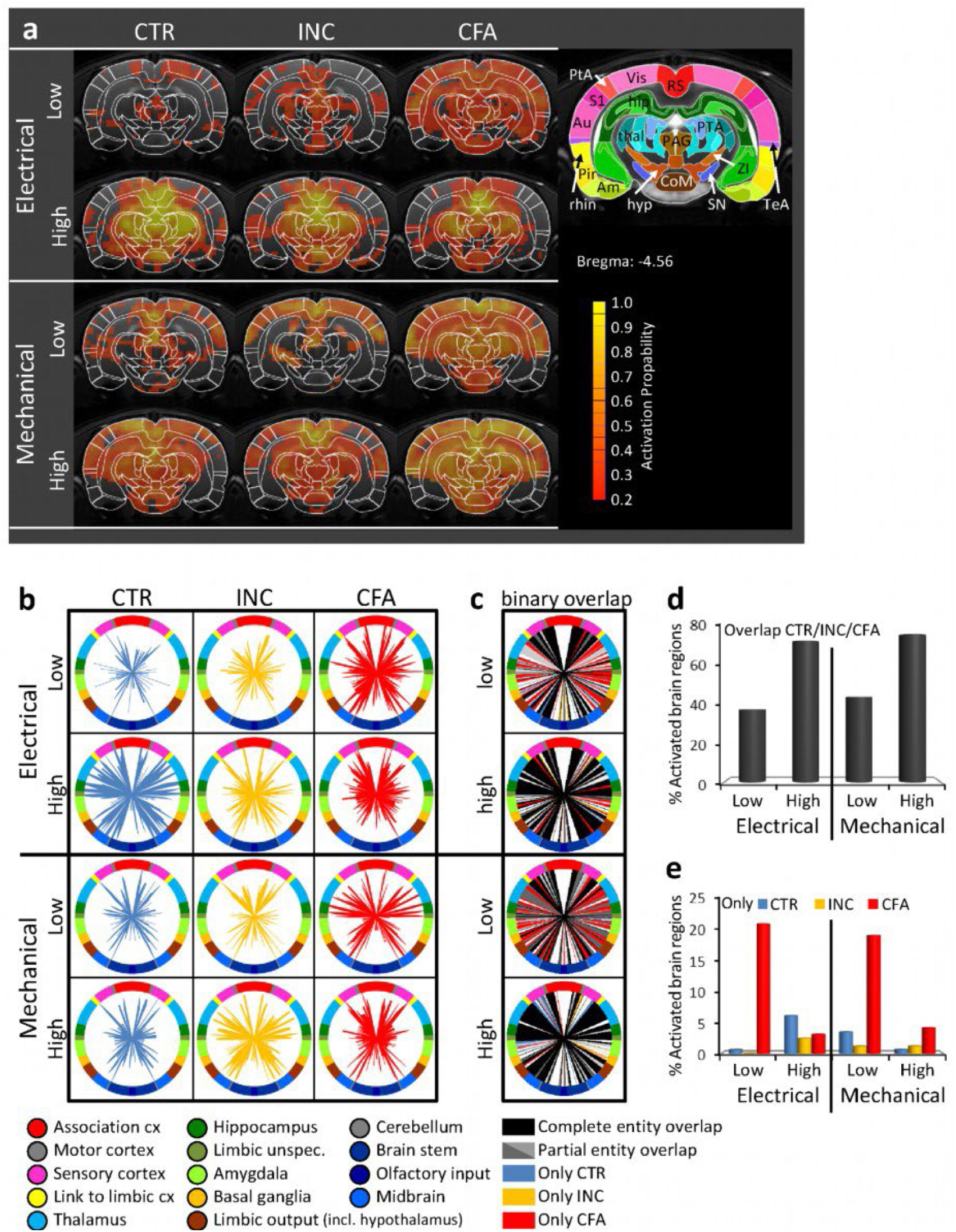
Activation probability of pain entities after electrical and mechanical stimulation with low (1mA / 50g) and high (5mA / 95g) intensities. **a)** Voxel-wise activation probability maps. **b)** Brain structure-specific activation probability. Length of spokes scales with the number of animals that showed activation in the respective region, referenced to the total number of animals in that group. **c)** Activation probability overlap (pain entity independent) of activated regions with at least 20% activation probability, encoded by the color of the spokes. **d)** Percentage of activated structures with complete overlap of all pain entities. **e)** Percentage of structures activated explicitly by only one entity. Am: amygala, Au: auditory cortex, CoM: corpora mammilaria, hip: hippocampus, hyp: hypothalamus, PAG: periaqueductal gray, Pir: piriform cortex, PtA: parietal association cortex, PTA: pretectal area, rhin: rhinale cortex, RS: retrosplenial cortex, S1: primary somatosensory cortex, SN: substancia nigra, TeA: temporal association cortex, thal: thalamus, Vis: visual cortex, ZI: zona incerta.

Consistent with earlier reports, bilateral BOLD activation was found, in pain-related regions for the ascending transmission (e.g., somatosensory cortex, thalamus, cingulate cortex, parabrachial nucleus, colliculi), descending modulation (e.g., periaqueductal gray, retrosplenial cortex), as well as for reward/motivation (e.g., prefrontal cortex) and affective/emotional components (e.g., amygdala, hippocampus, hypothalamus, anterior cingulate cortex) (Figure 2b,c) after unilateral peripheral stimulation (***Amirmohseni et al., 2016; Just et al., 2020; Sergeeva et al., 2015***). This basic activation pattern was primarily independent of entity (CTR, INC or CFA), modality (ES or MS), and intensity (low or high).

The activation probability increased with stimulus intensity for both modalities (Figure 2c, d). High-intensity stimulation resulted in the activation of up to 80% of all registered structures independent of the experimental group (black spokes in Figure 2c, and Fig 2d). Increased numbers of entity-specific activated brain structures were mainly observed during low stimulation intensity in CFA (Fig 2e), including midbrain structures (e.g., parabrachial nucleus, colliculi), motor cortex, hippocampus, and limbic output regions (Figure 2c). High-intensity ES resulted in reduced numbers of pain model-specific (INC and CFA) activated structures compared to CTR (Figure 2e). CTR animals specifically showed bilateral activation of pre- and infralimbic cortex and increased contralateral activation of the amygdala and sensory cortex (Fig 2c).

### Network size and density increases with stimulation intensity but varies with pain entity

Functional connectivity between activated structures revealed an enhancement of network density (only ES) and size (ES and MS) from low to high stimulation intensity in INC and CTR but not in CFA animals (Fig 3a, b). Except for high ES, INC showed networks with lower density and size (i.e., number of nodes) compared to CFA. For high ES, INC networks were denser and larger than CFA networks (Fig 3b). INC compared to CTR networks revealed the following: low ES resulted in large low-density networks for INC, whereas low MS resulted in smaller networks with higher density (Figure 3c). These findings suggest brain-wide specific differences in modality- and pain entity-specific cerebral processing.

**Figure 3:**
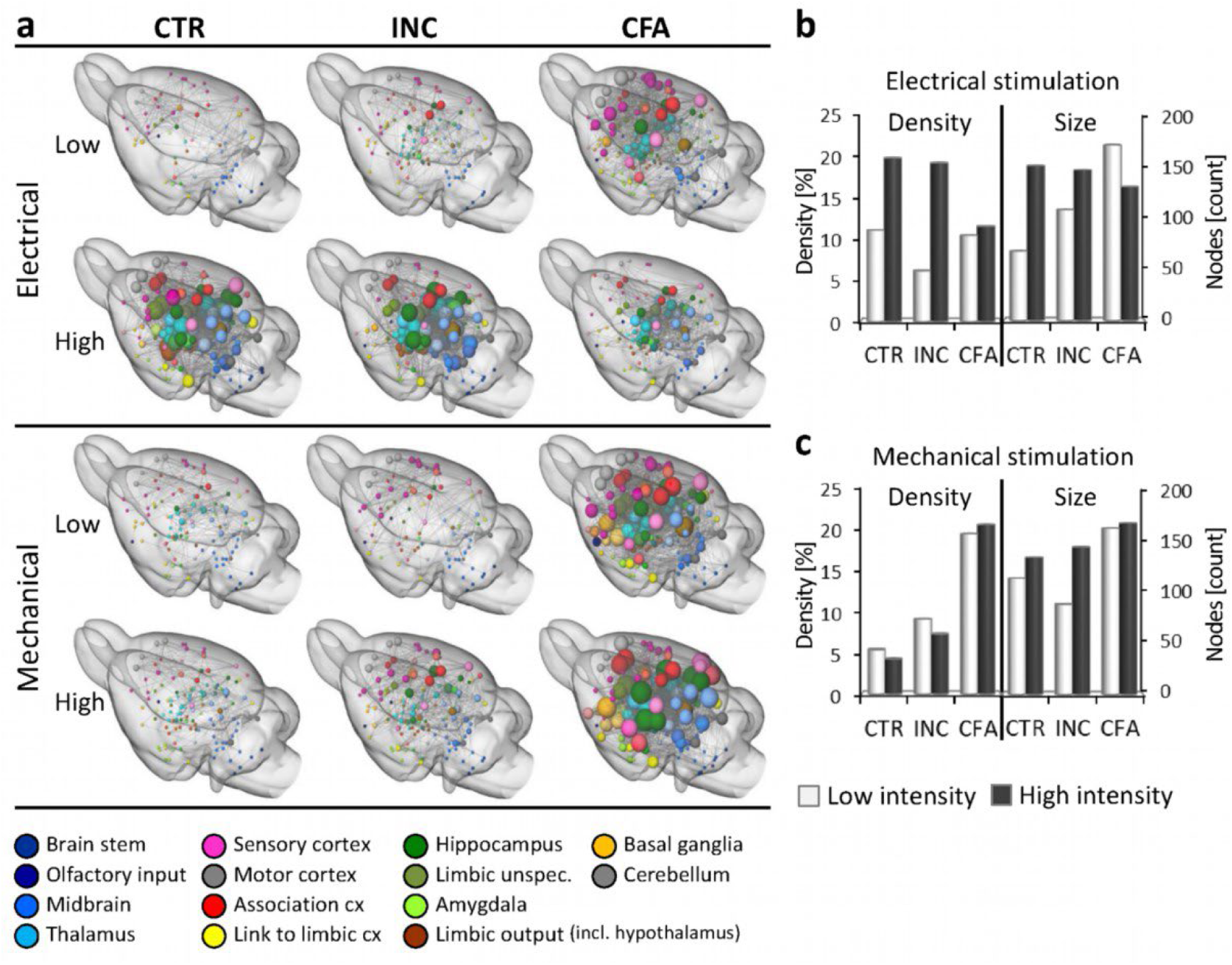
Networks of stimulus- and pain entity-specifically activated brain structures. **a)** Networks with anatomical node positions. All significant connections (determined by FDR with q=0.05) are displayed. Node size encodes degree, i.e., number of connections. Node colors represent anatomical groups of brain structures (see Supplementary Table 3). **b), c)** Network density (proportion of all possible connections) and network size (number of nodes) for electrical (**b**) and mechanical (**c**) stimulation. Only brain structures with an activation probability above 20% were taken into account.

### Small world index and numbers of communities depend on stimulation intensity

Next, efficiency of information flow in whole-brain networks was assessed by considering the small-world index (Fig 4a) and the number and structure of communities (Fig 4b,c). Both characterize the organization of brain networks with respect to segregation and integration. The small-world index σ is defined as the ratio of average clustering coefficient and average shortest path length of all network nodes, both compared to a random network of the same size (see Methods for details). A high σ is associated with increased efficiency of information flow. Since σ is dependent on network density, it was calculated for a range of densities between 3 and 20 %. In our data, in CTR and INC animals σ consistently decreased under high-intensity ES and MS conditions compared to low-intensity ES and MS, suggesting higher segregated networks during the processing of stimulation with high intensity (Figure 4a). In contrast in CFA animals σ increased for ES high-but decreased for MS high-intensities. A network density of 7% was used for all further graph-theoretical analyzes (close to the vertex of the high-intensity hyperbola curves, marked in Figure4a).

**Figure 4:**
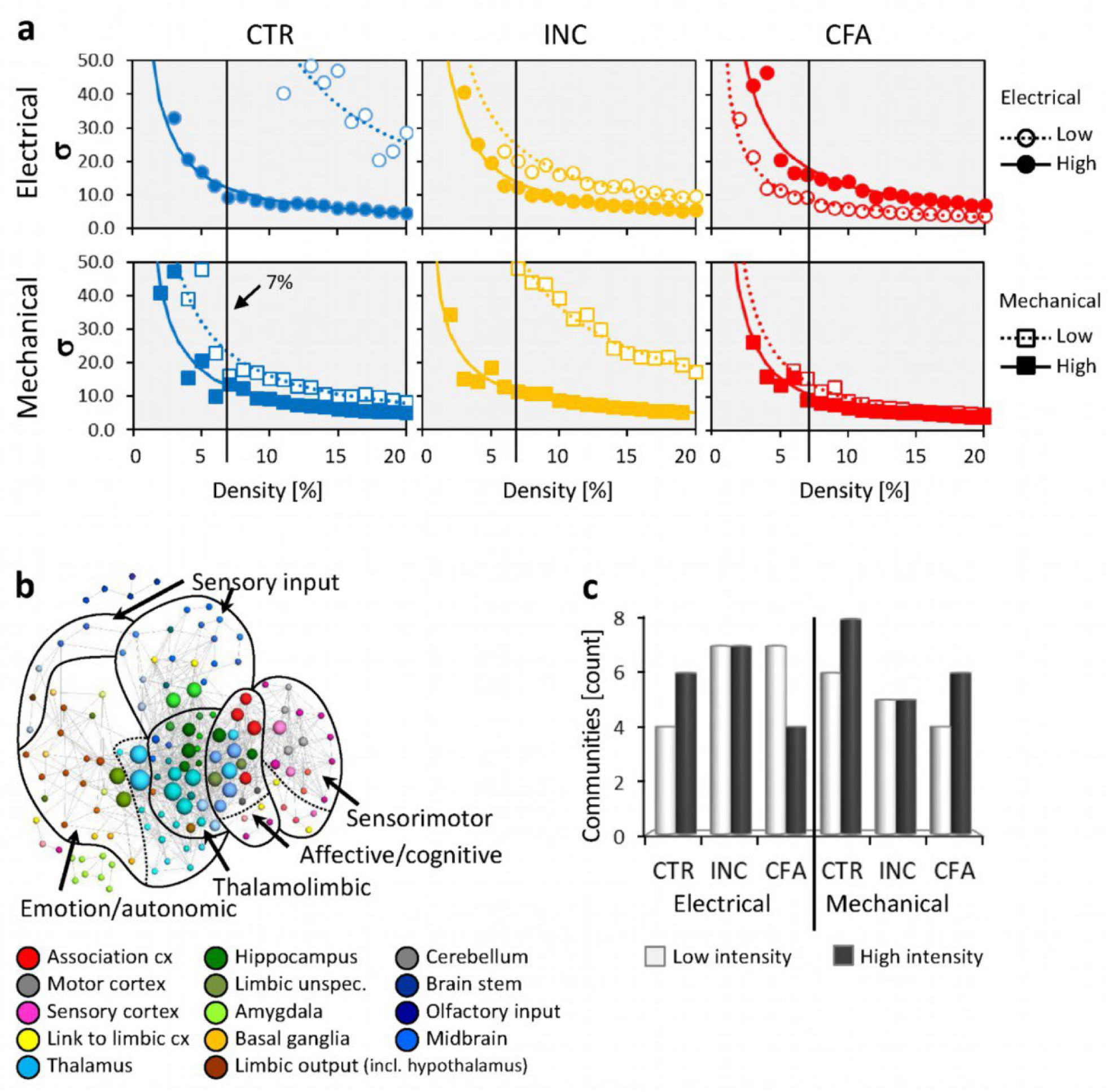
Global Parameters for segregation and integration of information flow: Small world index and communities. **a)** Small world index σ for average pain entity-specific networks over a density range from 3 to 20% for low and high ES and MS. The solid vertical line marks the density near the vertex of the high-intensity hyperbola curve. At this density (7% for all entities) the networks are fully connected but sparse enough not to be random. Therefore, graph-theoretical analysis was performed using a density of 7%. **b)** Communities and node distribution for CTR for the high ES average network (7% density). The node positions were determined using a force-based algorithm (***Kamada and Kawai, 1989***) suitable for visualizing communities. Node size encodes the respective degree. Node colors represent anatomical groups of brain structures (see Supplementary Table 3). **c)** Number of communities within average networks (7% density) of all pain entities after high and low ES and MS. Higher numbers of communities indicate higher segregation of functional brain units.

A set of nodes that are more densely connected within a community than with other nodes of the network characterizes the network community structure. Enhanced community numbers are a hint of enhanced segregation of functional brain units within the network. Network communities represent functional and anatomical entities of processing. For CTR and CFA, but not for INC, the count of intensity-specific network communities was related to the small-world index: CTR showed increased community numbers (4 to 6) and decreased small-word index with increased stimulation intensity. CFA (ES) showed decreased community numbers (7 to 4) and increased small-world index, respectively (Figure 4a, Supplementary Figure 2). In contrast, INC exhibited equal numbers if communities for both stimulus intensities but higher for ES (7) compared to MS (5).

CTR network communities associated with high-intensity ES resulted in the most comprehensive network in terms of involved brain structures and will therefore serve as the spatial node outline for further reference. It comprises the following functionally and anatomically related brain structures (Figure 4b):

1. Sensory input: brain regions of the midbrain and brainstem, dentate gyrus
2. Sensorimotor: sensory and motor cortex, parietal and temporal cortex
3. Thalamolimbic: (medial) thalamus, hippocampus, PAG
4. Affective/cognitive: cingulate and retrosplenial cortex, limbic cortex, (lateral) thalamus, cerebellum, habenulae, and colliculi
5. Emotion/autonomic: hypothalamus, basal ganglia, septum, amygdala, (ventral) thalamus

### Pain entity, stimulus modality, and intensity modulates connectivity strength

An indicator of information flow between brain regions is also represented by the strength of the functional connectivity (FC-strength) between nodes driven by the extent of neuronal synchrony. Modulations of FC-strength due to pain entity (high INC vs CTR and high CFA vs CTR), stimulation modality (ES vs MS) as well as intensity (high vs. low) were assessed by determining the largest interconnected component of connections with significantly (p< 0.05, uncorrected) altered FC-strength. The significance of the whole component compared to a random component of the same size (p_FWE_) was calculated using permutation testing (significance level p_FWE_<0.1, corrected) as previously reported (***Pradier et al., 2021***).

Comparing stimulus intensities for each stimulus- and pain modality revealed that increased ES intensity induced significant enhancement in FC-strength in INC and CTR pain entities (CTR and INC p_FWE_ <0.001, CFA p_FWE_=0.089), especially in the thalamolimbic and affective/cognitive community. In CTR animals, also structures of the emotion/autonomic and sensorimotor communities were modulated, which could not be observed in INC and CFA. Thus, for both pain models with experimentally induced hypersensitivity, the FC-strength of brain structures involved in the evaluative emotional and discriminative sensory aspect of central pain processing was independent of ES intensity (Figure 5a top).

**Figure 5:**
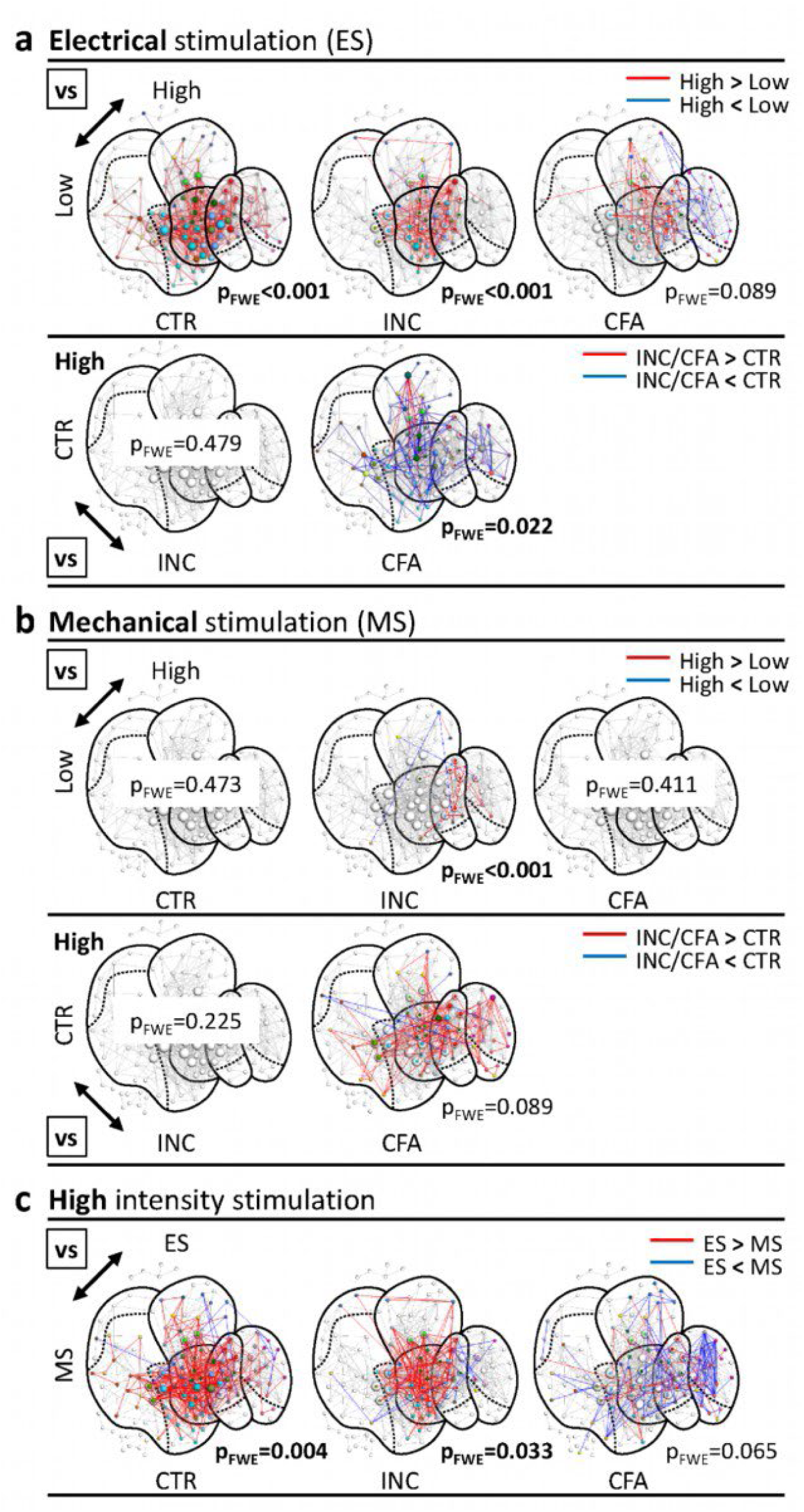
Impact of stimulus modality and intensity on pain entity. Changes in functional connectivities during **a)** electrical stimulation **b)** mechanical stimulation **c)** high intensity stimulation between ES and MS. Only significant components (NBS, pFWE<0.1) are shown for stimulation intensity (**a, b**, top), pain entity at high stimulation intensity (**a,b**, bottom), and high-intensity stimulation MS vs ES (**c**). Modulations imply for each connection significant (t-test, p<0.05, uncorrected) increase (red lines) or decrease (blue lines) of connectivity strength. Node positions were determined using the force-based Kamada-Kawai algorithm (***Kamada and Kawai, 1989***) derived from high-intensity ES CTR. Node size encodes the number of significantly modulated connections per node. pFWE: significance of modulated component, permutation corrected (NBS).

An increase of MS intensity resulted only for INC in a small but highly significant component (p_FWE_<0.001) of increased connections between cingulate (affective/cognitive community) and sensory cortex (sensorimotor community) (Figure 5b top). We could not detect any MS intensity-dependent FC-strength modulations in CTR, indicating that the MS at both intensities was processed similarly under control conditions.

When analyzing pain entities, INC at high stimulation intensity compared to CTR did not reveal significant components of modulated connections, neither for ES nor MS (Figure 5a,b, bottom). However, CFA showed a significant component of altered FC-strength (ES p_FWE_=0.022, MS p_FWE_=0.089) covering the same communities as the ES intensity-modulated component of the CTR group. This observation suggests that most connections relevant for pain processing were impaired in CFA, differentially modulated for stimulation modalities with decreasing FC-strength in ES (Figure 5a bottom) and increasing FC-strength in MS (Figure 5b bottom).

Additionally, we compared the FC strength between high-intensity ES and MS for all entities (Figure 5c). CTR showed a significant component (p_FWE_=0.004) of increased FC-strength in ES covering all communities, and CFA revealed a similar significant component (p_FWE_=0.065) of, again, decreased FC-strength in ES compared to MS. However, INC showed a smaller but significant component of increased FC-strength in the thalamolimbic and affective/cognitive community. Again, emotional/autonomic and sensorimotor pathways were not involved.

### Graph-theoretical node parameters indicate diminished entity-dependent network differences between stimulation modalities during a state of hypersensitivity

Detailed brain structure-specific modulations were further investigated by the following graph-theoretical network node parameters: strength and degree describing the node’s connectivity, clustering coefficient, and average path length as a measure of segregation or integration within the network, and finally, betweenness centrality and hypertext-induced topic selection (HITS) hub score, both characterizing the importance of the node for the network integrity and information flow. Since networks were compared at the same density (7%), alterations in the degree show a change in network size. All other parameters were normalized to the number of nodes and therefore were independent of network size.

To detect differences among sample means, we used three-factor ANOVA with factors stimulation modality (ES and MS), pain entity (CTR, INC and CFA) and brain structure (network nodes). Main effects in factors stimulation modality and entity show significant whole brain differences between factor group means. Stimulation modality with low intensity revealed a main effect only for connectivity strength (MS > ES) (Table 1, Supplementary Table 2 top). However, for high ES degree, average path length and betweenness centrality were significantly enhanced compared to MS (Table 1, Supplementary Table 2 bottom), indicating more extensive networks with more segregated node clusters after ES. The factor ‘pain entity’ was affected by strength, degree, and betweenness centrality at both intensities (Supplementary Table 2). ANOVA with follow-up Tukey HSD test revealed enhanced strength, degree, and betweenness centrality in CFA compared to factor groups INC and CTR for low intensity. However, high stimulation intensity leads to a decreased degree and betweenness centrality in both pain models INC and CFA compared to CTR, but enhanced connectivity strength for CFA compared to INC (Table 1).

**Table 1:** Post-hoc tests on ANOVA main effects for factor stimulation modality and factor pain model.

| | ANOVA Factor | | Strength | Degree | Clustering Coefficient<br>$\times 10^{-1}$ | Average path length | Betweenness centrality<br>$\times 10^{-3}$ |
| --- | --- | --- | --- | --- | --- | --- | --- |
| low stimulation intensity | stimulation modality | electrical | 1.45(0.05)<sup>a</sup> | no effect*<br>2.17(0.08) | no effect*<br>2.54(0.13) | no effect*<br>1.47(0.05) | no effect*<br>1.34(0.08) |
|  |  | mechanical | 1.73(0.06)<sup>b</sup> |  |  |  |  |
|  | pain entity | CTR | 1.25(0.06)<sup>a</sup> | 1.27(0.06)<sup>a</sup> | no effect*<br>2.54(0.16) | no effect*<br>1.47(0.06) | 1.07(0.09)<sup>a</sup> |
|  |  | INC | 1.33(0.07)<sup>a</sup> | 1.33(0.06)<sup>a</sup> |  |  | 0.98(0.08)<sup>a</sup> |
|  |  | CFA | 2.19(0.08)<sup>b</sup> | 3.91(0.15)<sup>b</sup> |  |  | 1.98(0.12)<sup>b</sup> |
| high stimulation intensity | stimulation modality | electrical | no effect*<br>2.19(0.06) | 3.94(0.10)<sup>a</sup> | no effect*<br>2.87(0.09) | 1.74(0.04)<sup>a</sup> | 2.41(0.10)<sup>a</sup> |
|  |  | mechanical |  | 3.29(0.10)<sup>b</sup> |  | 1.53(0.04)<sup>b</sup> | 1.74(0.13)<sup>b</sup> |
|  | pain entity | CTR | 2.12(0.07)<sup>ab</sup> | 3.97(0.12)<sup>a</sup> | no effect*<br>2.87(0.11) | no effect*<br>1.63(0.04) | 2.98(0.14)<sup>a</sup> |
|  |  | INC | 2.09(0.07)<sup>a</sup> | 3.40(0.12)<sup>b</sup> |  |  | 1.99(0.12)<sup>b</sup> |
|  |  | CFA | 2.36(0.08)<sup>b</sup> | 3.48(0.13)<sup>b</sup> |  |  | 1.88(0.16)<sup>b</sup> |

Data are the main effect of group averages with standard error in brackets. *: no significant main effect in three factors ANOVA (cf. Supplementary Table 2), values are averages and standard errors in brackets for all factor groups. a,b: Significant levels. Different letters show significant differences between group averages within each factor. ANOVA follow up Tukeys HSD, Bonferroni corrected, α=0.05. Note, that HITS hub score cannot show a main effect for pain entity and stimulation modality because we normalize the scores to the sum of all nodes within the network (i.e., the means are equal).

The factor brain structure always showed a significant main effect because the different brain structures intrinsically react differentially to external stimulation. Consequently, the interactions with other factors are more meaningful, indicating brain structure-specific modulations of the interacting factor. We found significant brain structure interactions only with stimulation modality for all parameters with high stimulation intensity and for low cluster coefficient, average path length, and HITS hub score (Supplementary Table 2). However, post-hoc t-test (FDR corrected, q=0.05) revealed only significant differences between high-intensity ES and MS in CTR. Here, ES enhanced parameter values predominantly in thalamic, limbic, and hippocampal brain structures (Figure 6). Additionally, the cingulum showed a bilaterally increased degree. It also elongated the average path length in the hypothalamus, zona incerta (limbic output), and cortical regions associated with emotion processing, i.e., contralateral (left) insula and ipsilateral (right) piriform cortex. However, the ipsilateral somatosensory cortex showed a reduced average path length showing higher integration into the functional network due to ES (Figure 6).

**Figure 6:**
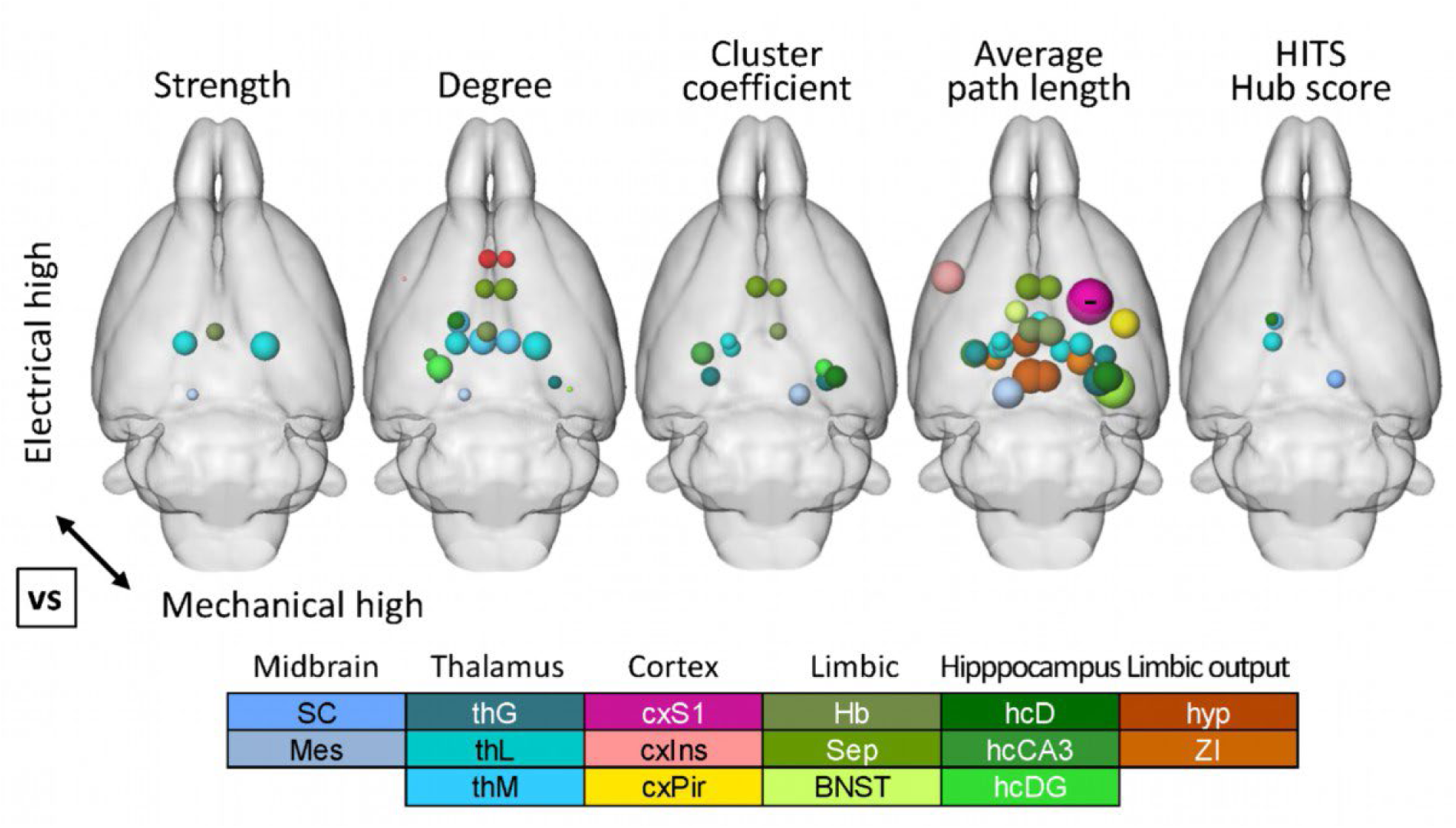
Significant modality-specific increase in node network parameters in “high-intensity” CTR ES vs MS networks. Displayed parameters showed significant interactions (p<0.05, corrected) with the factor brain structure in ANOVA of high-intensity stimulation (see Supplementary Table 2, bottom). Displayed nodes revealed significant enhancement using ANOVA post-hoc t-test, α>0.05, FDR corrected (q=0.1). Only the right cxS1 showed a significantly decreased average path length (marked with “-”). The significant ANOVA interaction between low-intensity stimulation modality and brain structure (Supplementary Table 2, top) could not be statistically verified. Node size represents the normalized difference between groups (ES-MS). BNST: bed nucleus of stria terminalis, cxIns: insular cortex, cxPir: piriform cortex, cxS1: primary somatosensory cortex, Hb: habenulae, hcCA3: CA3 field of the hippocampus, hcD: dorsal hippocampus, hcDG: dentate gyrus, hyp: hypothalamus, Mes: mesencephalon, SC: superior colliculus, Sep: septum, thG: geniculate thalamus, thL: lateral thalamus, thM: medial thalamus, ZI: zona incerta.

In summary, these findings show under CTR conditions, that ES and MS networks strongly differ, thereby indicating an ability of the brain to discriminate between both modalities. The additional aversive component for ES might explain the involvement of dominantly limbic and emotion-related cortical structures. These differences were less prominent for INC or CFA indicating a reduced modality discrimination under pathological pain conditions.

### Multivariate statistics reveal pain entities and stimulation modalities-specific biosignatures based on topological network parameters

Even though single graph-theoretical parameters may have relevance, the topology and information flow of a network can only be adequately characterized by simultaneously assessing all parameters, which is beyond classical univariate statistical testing. Therefore, we performed a linear discriminant analysis (LDA) using all the node parameter averages. LDA calculates linear combinations of parameters (linear discriminant dimensions, LD) with descending order of their class-separation contribution (later-on called separation) to classify groups. We show the projection scatter plots of classified brain structures in Supplementary Figure 3.

Relying on geometric means in the LD space of all brain structures per group (Figure 7), specifically for low stimulation intensity, LDA revealed a clear separation between CFA and both INC and CTR mainly along the first LD with 57.8% separation (Figure 7a top, gray arrow). Additionally, for both stimulation intensities graph-theoretical parameters could discriminate between ES and MS mainly along the second LD (low 21.2% and high 32.5% separation, Figure 7a,b bottom, solid black arrows), and selectively for high stimulation intensity along the first LD (45.4% variance, Figure 7B top, solid black arrow).

**Figure 7:**
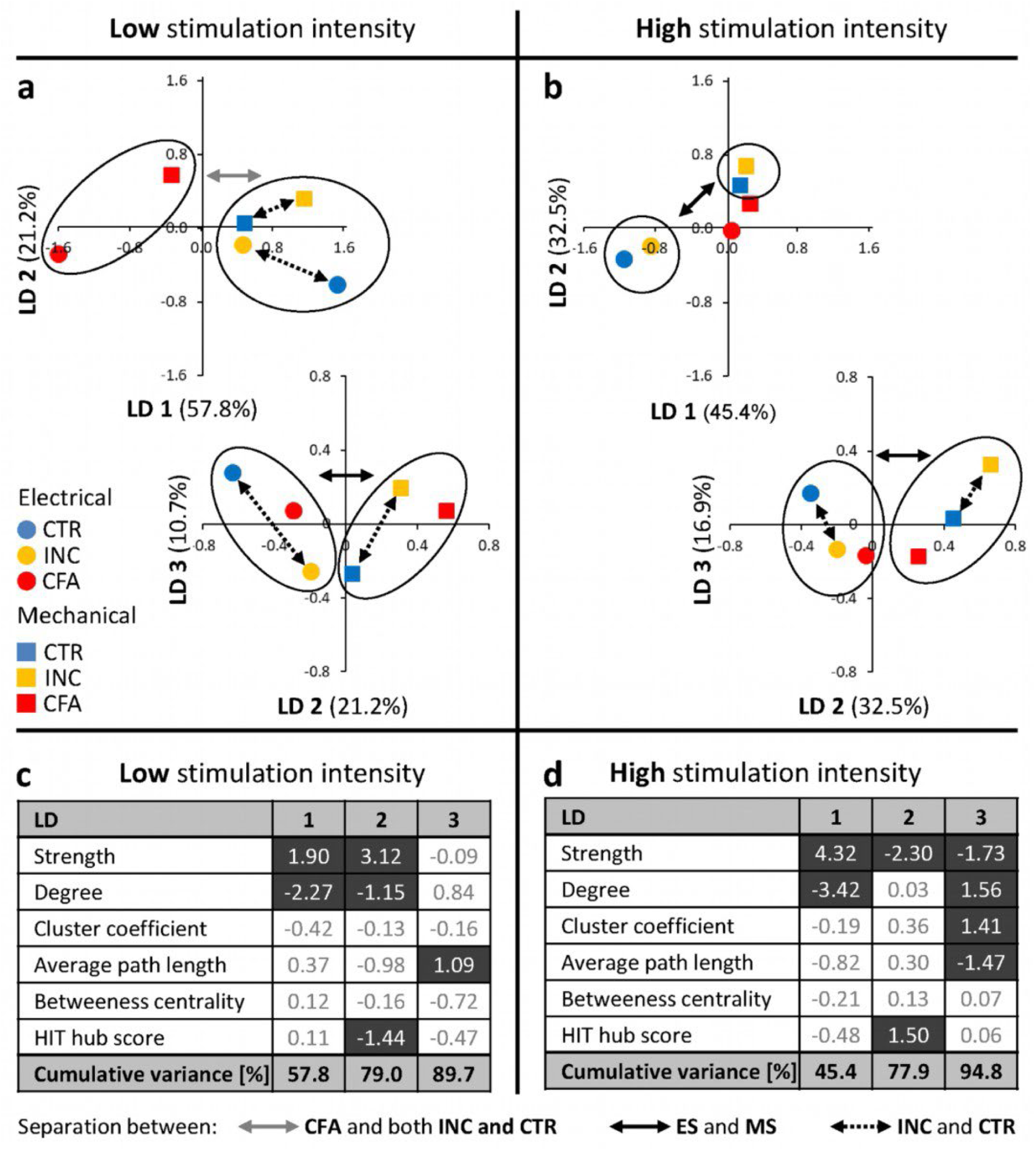
Biosignatures for separating of pain entity, stimulation modality, and intensity by graph-theoretical node parameters. Group separation was performed using Linear Discriminant Analysis (LDA). **a, b** group center of mass in Linear discriminant (LD) coordinates space for low (**a**) and high (**b**) stimulation intensities. Values in brackets show percentage of data variance that is represented by the respective LD. **c,d** Parameter loadings of the LD vectors resulting from the LDA of low (**c**) and high (**d**) stimulation intensity network node parameters. The higher the absolute value, the stronger the respective parameter’s contribution to the data variance assembled by the LD vector. We mark loadings above the absolute value of 1 with dark fields.

INC and CTR, but not CFA, could also be classified in ES and MS along the third LD (low 16.9% and high 10.7% separation, Figure 7a,b, bottom) and selectively for high stimulation intensity along the first LD (45.4% separation, Figure 7b top, black arrows). INC could be discriminated from CTR along the first two LDs only with low stimulation intensity. However, the third LD classified between INC and CTR regardless of the stimulation intensity (Figure 7, broken arrows). The contribution of the graph-theoretical node parameters to the first LD (i.e. the loading of the parameter on the LD, Figure 7 c,d) was dominated by connectivity strength and degree. For the second LD additionally the HITS hub score and for the third path length and only with high stimulation intensity also cluster coefficient gained classification importance.

Thus, the discrimination of CFA with low stimulation intensity was dominated mainly by quantitative aspects, such as strength and degree. However, ES and MS classification relied additionally on altered network topology (shown by high loadings of hub score, average path length, and clustering coefficient), showing altered information flow between brain structures. Focusing on INC, discrimination from CTR mainly included altered network topology, but only minor changes in quantitative network properties, especially with high stimulation intensity indicated by the small distance of group means along first and second LD.

### Specific brain structures discriminate between pain entities and stimulation modalities

Finally, we calculated the Mahalanobis distance between all pairs of corresponding brain structures in the LDA space between pain entities, modalities, and intensities (Supplementary Figure 4). The Mahalanobis distance describes the length but not the direction of the distance vector. The longer the Mahalanobis distance, the higher the respective brain structures’ contribution to the group discrimination. It defined most relevant brain structure pairs as those with distances longer than the mean of all pairs of two discriminated groups plus their standard deviation (horizontal lines in Figure 8, Supplementary Figure 4).

**Figure 8:**
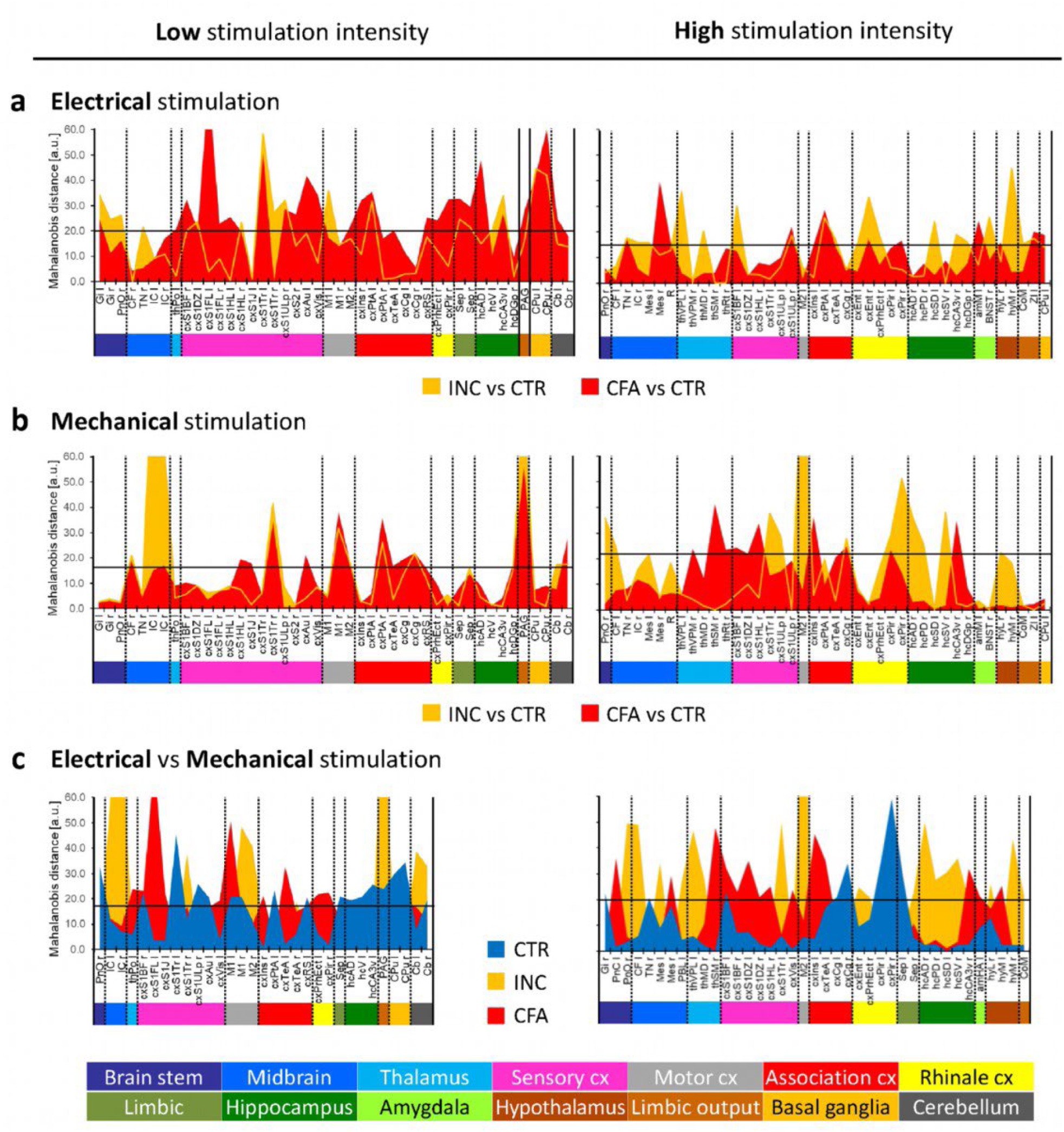
Mahalanobis distance between corresponding brain regions in LDA separated groups. Left: low stimulation intensity. Right: high stimulation intensity. It displayed only distances of brain structures that exceed (in at least one comparison per stimulation intensity) the average distance + standard deviation, which is shown by the horizontal line in each diagram. Brain structures we pooled into anatomical groups, marked by colored bars below the x-axis. **a,b** Mahalanobis distance between CTR and pain models (INC, CFA) after electrical (**a**) or mechanical (**b**) stimulation. It calculated separately the average distance and standard deviation for each stimulation modality, the chosen set of brain structures comprised both modalities. **c** Mahalanobis distance between ES and MS. We calculated the average distance and standard deviation over all pain entities, resulting in one set of brain structure per stimulation intensity.

For ES, a separation between CTR and pain models, indicated by a high Mahalanobis distance, was dominated by CFA for low and INC for high stimulation intensity. Brain structures that discriminate most between CTR and both pain models at low ES stimulation intensity were S1 (mainly trunk) and the caudate putamen. Specifically, CFA showed long distances of the contralateral fore and hind limb fields of S1 and limbic areas (piriform cortex, septum, dorsal hippocampus) to the corresponding CTR brain structures (Figure 8a left). At high ES, especially the lateral thalamus, the entorhinal cortex, and the hypothalamus showed larger distances between INC and CTR than between CFA and CTR (Figure 8a right).

For MS, low intensity led to comparable brain region distances to CTR for both pain entities. Note that the distance vectors may have different directions despite the same length. These structures comprised mainly S1 (trunk field), motor cortex, parietal and temporal association cortex, cingulum, and the longest distance, the PAG. Solely, the inferior colliculi showed a much longer distance of INC to CTR (Figure 8b left). For high MS stimulation, INC and CFA discrimination from CTR involved different brain structures. CFA was dominated by long distances to CTR of thalamus and S1, whereas INC showed the longest distances to CTR for contralateral M2, piriform cortex, and hippocampus (Figure 8b right).

The distances between ES and MS at low stimulation intensity were dominated for INC by the inferior colliculus and the PAG, for CFA by the left S1 forelimb, motor cortex, and temporal association cortex, and for CTR by the left S1 trunk field, pons, and caudate putamen. With high-intensity stimulation, additional structures, especially thalamus and limbic systems, contributed to the distance between ES and MS. Whereas discrimination of ES and MS in the CFA group was still dominated by long distances of the S1 and temporal association cortex, the INC group also involved pons, thalamus, hippocampus, and hypothalamus. In the CTR group, we observed the most extended distances for the piriform and cingulate cortex.

### Summary of local network parameters

To summarize our main findings, we combined averages of normalized differences of significantly changed node parameters (Figure 9). We focused on clinically relevant low-intensity modalities because these more closely resemble a state of clinically relevant hyperalgesia (for summary of high-intensity modalities see Figure 9). After INC and CFA, we observed increased network modulation following ES and MS in affective/cognitive, cognitive-motor coordination (motor cortex) and autonomic (hypothalamus) network components. Furthermore, after CFA, we observed globally increased network modulation during MS and ES compared to CTR (Figure 9, right column), while after INC, we noted reduced network modulation in the brainstem (ES and MS), thalamus (MS), somatosensory cortex (ES), and the caudate putamen (ES and MS), which are part of the ascending sensory/discriminative pain pathway and desending subcortical motor coordination (Figure 9, left column).

**Figure 9:**
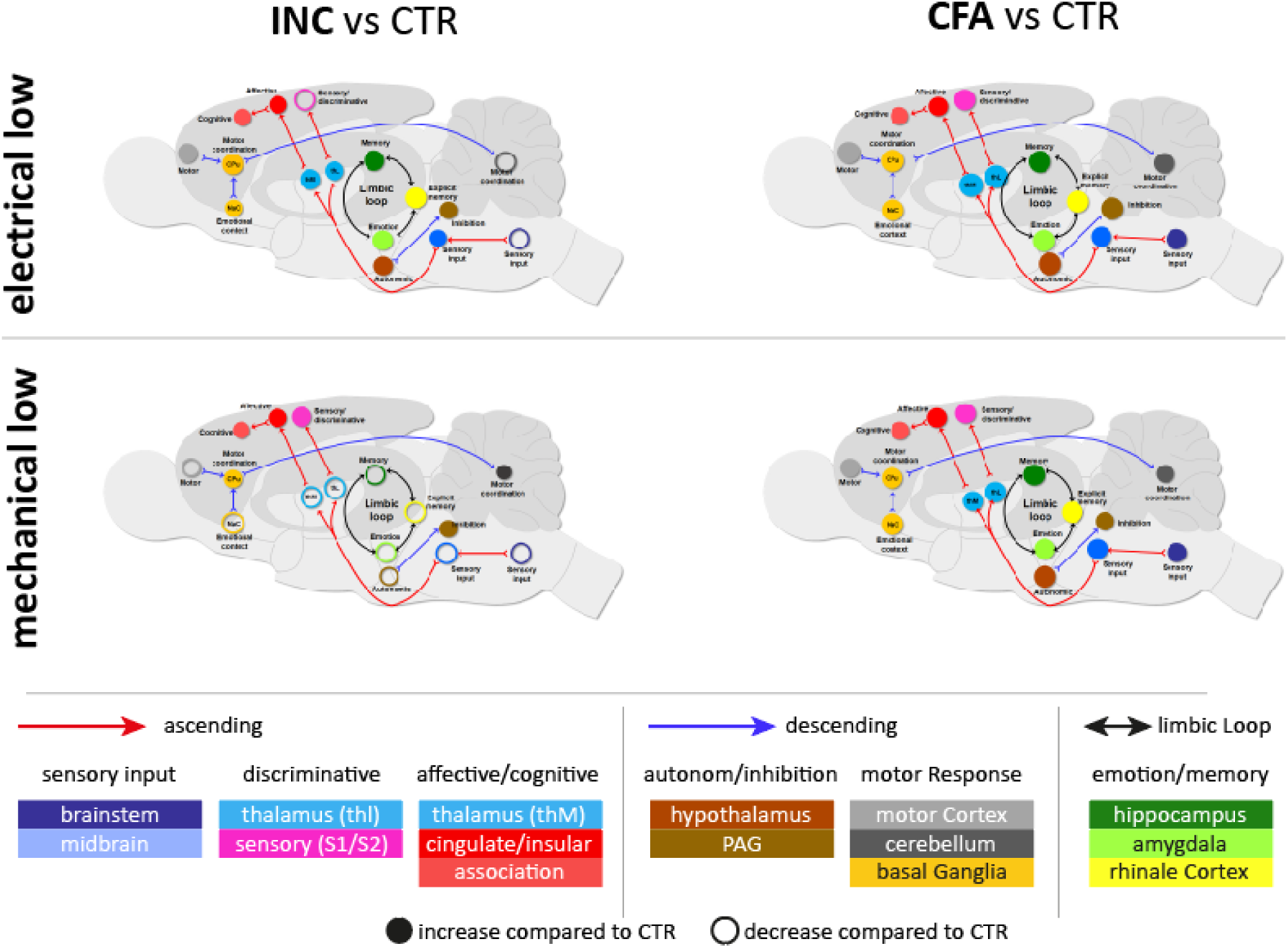
Summarized network modulation of pain entities compared to control. Average normalized difference to control of complete network node degree (cf. Figure 3) and significantly modulated node parameter revealed by the graph-theoretical and LDA analyzes performed in this study. These are the total of the absolute values of decreased (−1) and increased (+1) connections per node in significantly modulated components (cf. Figure 5), those node parameters that show a significant interaction of ANOVA factors pain entity, pain modality and brain structure (average path length and hub score for low, and strength for high intensity stimulation, cf. Supplementary Table 2), and the high Mahalanobis distances to CTR revealed by LDA (cf. Figure 8). We inverted the range of the average path length values to ensure that all parameter values point towards increased network integration. The Mahalanobis distance has no direction and was therefore only integrated into the calculation of the absolute difference to control. Node size codes for the absolute average normalized difference, filled nodes show enhancement, and empty nodes indicate a reduction of network integration compared to control. Visualization focuses on ascending (red line) and descending (blue line) pain pathways characterizing pain processing in the brain. CPu: caudate Putamen, NaC: nucleus accumbens, thL: lateral thalamus, thM: medial thalamus.

## Discussion

Specific neuroimaging signatures are an important step toward identifying novel pain biomarkers that can be applied to develop and validate novel therapeutic options, which ultimately can be translated to the clinics (***Tracey, 2021***). Because different pain entities diverge regarding the mechanism to induce or maintain pain-related outcomes, a pain-entity-specific approach is necessary. Yet, the challenges of identifying neuroimaging signatures at a pain entity- and modality-specific level are manifold. Therefore, we developed a comprehensive framework for brain network analysis in disease-specific pain models combining functional MRI with graph-theory and data classification by linear discriminant analysis. This enabled us to expand our knowledge of stimulus (mechanical vs. electrical) modality processing under incisional (INC) and pathogen-induced inflammatory (CFA) pain entities compared to acute pain control conditions. In short, graph-theoretical analyses revealed distinct Network Signatures of Pain Hypersensitivity (NSPH) for INC and CFA, resulting in impaired discrimination of stimulus modalities in both pain models compared to control conditions (CTR).

### Different modalities result in distinct network signatures in CTR condition

Preclinical pain studies in animals predominantly use peripheral stimulation to measure hypersensitivity as a proxy of the pain symptoms of hyperalgesia and allodynia in humans. Hypersensitivity to mechanical pain is a prominent symptom in human pain disorders, yet the investigation of its mechanisms is virtually absent in preclinical fMRI animal studies. To increase the translational value of small animal fMRI, our newly developed mechanical stimulator for rats (***Amirmohseni et al., 2016; Just et al., 2020***) allowed us to compare the activation of mechanosensitive Aβ, Aδ and C-fibers (***Zimmermann et al., 2009***) with the less selective activation of peripheral fibers through electrical stimulation and to investigate their effects on pain (hyper-) sensitivity. Since both MS intensities used here are suprathreshold for mechano-sensitive fibers (***Zimmermann et al., 2009***), we conclude that similar network processing resulted from peripheral inputs composed of comparable fiber populations. Interestingly, we observed strong differences in ES between both intensities: employing a similar stimulus protocol, previous studies have shown that 1 mA activated non-nociceptive low-threshold Aβ and Aα fibers, but not polymodal C-fiber nociceptor (***Le Bars et al., 1979***). Stimulus intensities exceeding 1.5 mA resulted in pain-related behavioral responses (***Zhao et al., 2014***) and additional activation of mechanosensitive Aδ and polymodal (thermo- and mechano-sensitive) C-fibers (***Koga et al., 2005***). We, therefore, suggest that the bulk increase in changes related to FC and local network parameters with high (5 mA) ES intensity resulted from the additional activation of heat/cold and mechanical nociceptor, that leads to an overall integration of brain regions involved in the processing of pain and aversion without any specificity to pain-related symptoms evoked by stimulus modalities like pricking the skin or heat.

The strong difference between MS and ES in network processing at the FC- and local network level in CTR rats suggested an ability of the brain to discriminate between modalities and involved changes along the ascending and descending pain pathways. Brain regions that discriminated best between ES and MS at low intensities were S1 trunk, M1, hippocampus, and striatum, while for discrimination of ES and MS at high intensity, the piriform and cingulate cortex were the most different. These results parallel findings from a recent study in humans that compared aversive processing of mechanical and heat stimulus modalities (***Čeko et al., 2022***); here, the S1 region was found as the modality-selective region for mechanical pain, while M1, cingulate cortex and the striatum was selective for heat pain. However, in that study, the anterior and medial insular cortex also predicted the averseness of noxious heat stimuli. This was not the case in our study and hence could reflect species differences under naïve conditions or averseness differences between noxious heat and mechanical stimulation. Other modality differences were likely related to encoding positive or negative stimulation values (***Han et al., 2020***) in the dorsal hippocampus and noxious stimulus intensity in the piriform cortex (***Lötsch et al., 2020***). Together, the modality-specific differences uncovered by our comprehensive analysis framework establish key brain regions important for processing sensory discrimination and aversive aspects of pain, as well as regions involved in the multisensory integration of this information.

### Global differences in entity-specific network processing despite similar behavior

The two pain-models, surgical incision, and pathogen-induced inflammation, both led to peripheral and central sensitization and resulted in mechanical hypersensitivity as a prominent symptom. This is consistent with our recent reports (***Segelcke et al., 2021b; Segelcke et al., 2021a***), in which we compared both pain models in a multimodal behavioral test battery: heat- and mechanically evoked as well as non-evoked pain-related responses, and gait parameters did not differ 24h after induction in both models. This was also true for peripheral macroscopic parameters of the inflammatory response (edema and hind paw skin temperature (***Segelcke et al., 2021b***). However, strong differences existed regarding the mechanisms to induce hypersensitivity in both pain states. The incision, a model for local tissue injury-based peripheral sensitization, mainly recruits immune cells and leads to enhance cytokine and proinflammatory mediator concentration locally (***Pogatzki-Zahn, 2017; Segelcke et al., 2021b; Segelcke et al., 2021a***). It causes inflammation that affects skin, muscles, and internal organs, but not the brain directly, although this has not been precisely investigated yet (***Ji et al., 2018; Matsuda et al., 2019***). In contrast, peripherally injected CFA models infection-induced inflammation and induces longer-lasting immune responses that lead to a more general sensitization via various cytokines and inflammatory mediators (***Huang et al., 2020; Julius and Basbaum, 2001; Matsuda et al., 2019; Segelcke et al., 2021b; Segelcke et al., 2021a; Sergeeva et al., 2015***). Moreover, CFA induces a so-called neurogenic inflammation and promotes inflammation in multiple brain regions, including midbrain, thalamic, and forebrain regions (***Griton and Konsman, 2018; Raghavendra et al., 2004; Tan et al., 2019; Torsney, 2019***) probably through priming of the choroid plexus (***Shrestha et al., 2014***). In line with this, our data - specifically at low intensity stimulation - show globally greater BOLD activation, FC, and network modulation in CFA animals compared to CTR, while changes in INC animals were instead at an intermediate level between CFA and CTR after stimulation. Therefore, our data suggest that CFA treatment results in a stronger central (cerebral) sensitization associated with an increased neuro-inflammatory component compared to the INC model.

### Common and distinct features of pain entity-specific hypersensitivity neuroimaging signatures

A hallmark of pain hypersensitivity in rodent studies is the increased nocifensive behavior following noxious or innocuous stimulation compared to the control condition. To avoid ceiling effects, we therefore suggest that hypersensitivity-related signatures are best studied at low stimulus intensities. Indeed, the regions that discriminated INC and CFA best from CTR conditions showed a striking overlap with those established to be important for modality-specific processing and included S1 trunc, M1, striatum, and cingulate cortex. However, the modality-specificity we observed under CTR conditions no longer exists under CFA/INC conditions. Importantly, it has been demonstrated that under conditions of inflammatory (CFA) and chronic pain, the S1 cortex shifts from sensory discrimination to the mediation of aversive behaviors (***Jin et al., 2020; Singh et al., 2020***) through inputs to the anterior cingulate cortex and the dorsolateral striatum. Given the different roles of these regions in CTR compared to CFA and (putatively) INC conditions, this suggests that reduced sensory discrimination and increased aversion characterize NSPH. Additionally, we identified the PAG and cuneiform nucleus (under MS) as well as the hippocampal CA3 and the septal area (under ES) as key regions for hypersensitivity. Interestingly, PAG/CfN (***Carlson et al., 2005***) and CA3/septal regions (***Kim et al., 2014; Luo et al., 2011; Schvarcz, 1993***) have been reported to mediate descending pain modulation or providing pain relief, respectively. This further indicates that NSPH have additional – and possibly modality-specific - pain modulating components, probably to compensate the increased nociceptive and aversive input during hypersensitivity. Furthermore, we also detected the parietal cortex as part of NSPH, which could be involved in the discrimination between nociceptive and salient stimuli (***Horing et al., 2019***) as well as stimulus localization (***Porro et al., 2007***).

Importantly, the direction of network modulation was not always the same in CFA and INC compared to CTR, and it partly decreased components of the ascending input in INC. This suggests that both pain models mainly differ in their descending modulation of sensory input. Moreover, we observed additional activation of the insular cortex and the anterio-dorsal hippocampus under CFA conditions. Both regions have strong connections to pain processing and mediate nociceptive hypersensitivity and supposedly link sensory and affective-motivational networks (***Lu et al., 2016***), (***Barthas et al., 2015***) as well as pain relief (***Wei et al., 2021***), respectively.

As it is obvious by the many human neuroimaging studies referenced above, our established analysis fMRI framework can very directly be translated to human patients to characterize their pain symptoms and relief at a very fine, brain structure specific way as exemplified in Hess et al. (***Hess et al., 2011***).

### Strengths and limitations of the analysis framework for hypersensitive brain networks

The graph-theoretical analysis used in this work focuses on functional relationships between regions, thereby reflecting the dynamics of underlying information processing. In addition to the static activation patterns (i.e. BOLD), graph-theoretical parameters characterize the information flow in much more detail following noxious and innoxious stimulation. Therefore, we could detect specific network signatures characterizing different pain entities and pain modalities, which was not able relying on activation patterns alone – thereby illustrating the power of graph-theoretical analysis.

Relying on Hebb’s theorem “fire together wire together” (***Hebb, 1949; Keysers and Gazzola, 2014***), graph-theoretical analysis of neurological data relies solely on the mathematical similarity of time courses – in this work, the Pearson’s correlation coefficient of regional time courses derived from task-related fMRI. This implicates several methodological issues that have to be addressed during analysis.

First, the correlation coefficient and its statistical significance are influenced by the number of sample points (determined by TR) reflecting the time course. The TR of 1s and 600 time points per scan used here were sufficient to obtain reliable results; however, these issues should be taken into account when comparing results from different studies.

Second, physiological and thermal noise also influences time courses. Since noise is unspecifically distributed across the whole brain, it enhances correlation strength but reduces regional specificity. Besides optimal data acquisition and appropriate preprocessing global noise can be reduced by global signal regression. Concurrently global signal regression also eliminates the signal fluctuation induced by the stimulation leaving time course fluctuation dominantly accounting for functional connectivity instead for stimulation response. However, global signal regression is discussed controversially since it artificially induces high amounts of negative correlations (***Murphy et al., 2009***). On the other hand, it enhances the specificity of detected functional connections (***Fox et al., 2009; Liang et al., 2012***), especially when focusing only on positive correlations.

One core characteristic of graph-theoretical network analysis is a custom-defined density that distinguishes organized from random graphs. If every node is connected with every other node, which represents the maximum density of a graph, then this graph is equal to a fully connected random graph. To identify differences in network organizations/topologies, the densities of the investigated networks have to be the same. Therefore, defining a network density is a crucial step in the graph-theoretical analysis. Usually, this is done arbitrarily, often following the rule that the network has to be sparse but fully connected. However, this rule ignores that some brain regions, although statistically activated, play a minor role in a neural network accompanying task processing. Thus, a fully connected network might be too dense to gain meaningful information. In this work, we introduced a more objective density definition relying on the hyperbolic curve of the small world index, a measure for network organization compared to random networks over a wide range of network densities. We used the next highest density to the vertex of the fitted hyperbola, representing an objective and network-specific point of balance between high connectivity and high organization level.

Statistical approaches to evaluate network differences between groups, in this work, pathological pain entities compared to pain control entity, have to deal with two major features: the great amount of single tests and the great amount of parameters describing the network and the role of individual nodes (i.e., brain structures) within the network. Since classical approaches to correct for multiple comparisons are usually too conservative in detecting any group differences in complex networks, NBS was developed (***Zalesky et al., 2010***). However, NBS is a weak statistic, revealing a component of contiguous suprathreshold edges whose statistical significance is determined using non-parametric permutation testing. The importance of a single edge within this component cannot be assessed. The integration of the various graph-theoretic parameters and derived features for group classification can, in principle, also be done with machine-learning approaches (***Lamichhane et al., 2021***). However, the small number of animals always hampers the application of machine-learning approaches. Therefore, we used the classical linear discriminant analysis, which is much more robust to small sample sizes.

Finally, the anesthesia used here for animal fMRI studies is always a confounder and in particular hampers the full translational power of fMRI. However, the anesthesia was identical for all pain models and types of stimulations used, allowing direct comparisons. We are not aware of any specific hypothesis that a given anesthesia impacts selectively either experimental condition. In contrast, it is overall demonstrated in many studies, that the principal brain activation between anesthetized animals and awake humans induced by nociception is directly comparable.

## Conclusion

Our new analysis framework allowed us to extract specific features for pain modality processing in rats that well translate to those previously detected in humans under naïve conditions and include S1, M1, CPu, HC, piriform and cingulate cortex. Further, we could detect pain entity-specific network signatures for incision, inflammation and acute pain, that share decreased modality discrimination, but increased aversion processing and descending pain modulation. These signatures included PAG, Cf, CA3 and septal area, as well as the parietal cortex. Our data further suggests that pain entities modulate sensory input by different Pain Hypersensitivity-Specific Network Signatures which, as such, can be directly translated to human patients suffering, e.g. from postoperative hypersensitivities.

## Methods (max. 3000 words: 1969 words)

### Animals

We used fMRI raw data from 21 male Sprague-Dawley rats (SD, 270-350 g) obtained from our previous study (***Amirmohseni et al., 2016***) to investigate functional brain networks for different pain entities (models for incisional pain: INC and infection-induced adjuvant/pathogens-inflammation pain: CFA), modalities (electrical: ES and mechanical: MS stimulation), and intensities (low and high). The State Agency reviewed all animal experiments for Nature, Environment and Consumer Protection North Rhine-Westphalia (LANUV), and are in accordance with the ethical guidelines for investigation of experimental pain in conscious animals.

### Pain entities

Briefly, the incisional pain model (INC) was made by an incision on the plantar aspect, and the infection-induced inflammation pain model (CFA) was induced by intraplantar injection of 150 µl of complete Freund’s adjuvant at the right hindpaw. We performed all interventions under general isoflurane anesthesia. Sham (CTR) animals received the same anesthetic regimen to compensate for possible anesthesia effects and serve as a control for both groups. All rats were allowed to recover for 24 h in their home cages before fMRI measurement started.

### Behavioral tests

One day before (baseline) and 24 h after model induction, the rats were subjected to a behavioral test for their mechanical pain threshold (von Frey). Rats were placed individually on an elevated plastic mesh floor (5 x 5 mm) covered with a clear plastic cage top (21 × 27 × 15 cm^3^). After 30 min of acclimatization, calibrated Semmes-Weinstein von Frey filaments were applied from underneath the mesh to the plantar aspect of the proximal part of the heel of one paw. The mechanical force via the filament was increased, starting with 1.35 g until a withdrawal response occurred or the cut-off of 60 g was reached. We considered the median of three repetitive tests per rat as the paw withdrawal threshold. Significant differences between baseline and 24 h post model induction were determined using the non-parametric Friedmann and Dunnett test.

### fMRI acquisition

Rats were fixed in the prone position on the fMRI bed under general isoflurane anesthesia one-day post pain entity induction and 3h after the behavioral test. Vital parameters (body temperature, respiration/heart rate, pCO_2_) were monitored during the entire acquisition phase and were maintained in the normal range. fMRI measurements were conducted on a 9.4 T Bruker Biospec 94/20 small animal scanner (Bruker Biospin GmbH, Ettlingen, Germany) using a single loop surface coil (Bruker). Adjustments were conducted and anatomical images (RARE: TR/TE = 4000/12.5 ms; RARE factor = 8; FOV = 28 × 26 mm2; 256 × 256; 16 slices; slice thickness = 1.2 mm; 4 averages) were acquired, followed by shimming using MAPSHIM (Bruker) over the entire brain. The right hindpaw was secured and accessible for ES and MS. The anesthesia regime was switched to medetomidine to avoid isoflurane effect on BOLD-signal after. The stimulation protocol was a block design comprising 10s OFF-10s ON-10s OFF intervals repeated 20 times (total 600 s). Electrical stimulations (square wave pulses, DS5, Digitimer, Welwyn Garden City, UK) were applied subcutaneously between the second and fourth digit of the right hindpaw with two intensities (1mA:low and 5mA: high at a frequency of 9 Hz and with 2ms pulse duration). Mechanical stimulation of the plantar aspect of the right hindpaw was performed using an in-house made MR-compatible pneumatically-controlled device (***Just et al., 2020***) with a von Frey filament (0.85 mm diameter, yielding a contact area of 0.56 mm^2^) and two different intensities (50g (low) and 95g (high) at a frequency of 1Hz with 0.5s duration).

### fMRI-data preprocessing

Preprocessing of fMRI raw data included inter-slice time and motion correction (3D with trilinear/sinc interpolation), linear detrending (high pass filter, cut off 0.015 Hz), and spatial (Gaussian, FWHW 1 mm) and temporal (Gaussian, FWHW 3 sec) smoothing. Subsequently, a General Linear Model Analysis (GLM) was performed with stimulation intensities as separate predictors. Following FDR correction (q=0.05), the corresponding threshold was applied, thereby obtaining statistical parametric maps (SPM) to identify significantly activated voxels per animal and stimulation condition.

### Activation probability

For SPM group comparisons, the first volumes of all EPI time courses were registered affine, with 5 degrees of freedom to the same template. This template was an EPI volume of one selected sham animal. The resulting transformation matrix was used to register the thresholded and binarized SPMs, and group averages were obtained for each pain entity, stimulation modality, and stimulus intensity. These group averages correspond to activation probability maps, indicating spatially (voxel-wise) the proportion of animals with significant activation.

An in-house developed digital brain atlas based on the Paxinos rat brain atlas (***Paxinos and Watson, 2007***) was manually registered (affine, 5 degrees of freedom) slice-wise to the average first EPI volume. Backward transformation of the resulting brain atlas using the transformation matrix of the EPI volume registration described above leads to individual atlas labeling in its native space. The atlas comprised 202 ROIs representing specific brain structures (Supplementary Table 3).

Multiplying these atlas labels with the corresponding binarized SPMs resulted in defined activated voxels per brain structure, stimulus condition and animal. Analogous to the voxel-wise activation probability, a region-wise activation probability was calculated by determining the proportion of rats within each group that showed at least one activated voxel in a specific structure. In order to minimize outlier effects, for further analysis, only structures with an activation probability greater than 20% (i.e., at least two animals showed structure-specific activation) were considered.

### Functional connectivity

For each animal and stimulation condition, the average time courses of all activated voxel per region were cross-correlated, resulting in symmetric correlation matrices. To obtain group-specific networks, first, we averaged separately these correlation matrices for each pain model, stimulation modality, and stimulation intensity. For each average group matrix, significant correlation coefficients (Pearson’s r) were determined and corrected for multiple comparisons using FDR (q=0.05). On average, significant r-values were higher than 0.113 ± 0.008 (Supplementary Table 1). Second, these thresholded average group matrices were translated into networks with nodes and edges, representing the activated brain structures and the functional connections between pairs of brain structures. The weight of each edge corresponded to its correlation coefficient r and showed the strength of the given functional connection.

Since the networks of the different stimulation conditions had different numbers of activated structures and different numbers of significant functional connections the resulting group-specific networks had different sizes. In general, graph topology and derived graph-theoretical parameters are dependent on network density. Therefore, all group networks were adjusted to the same density, with the strongest 7% of all connections. This density allows graphs that are fully connected but sparse enough to show a topological network organization distinct from a random network.

### Graph-theoretical network analysis

Typical characteristics of complex networks are organization in communities and a high small-world index (***Watts and Strogatz, 1998***). Network communities are sets of nodes that are more densely connected internally than with other nodes within the network. For each network, communities were detected using the algorithm proposed by Blondel (***Blondel et al., 2008***) and visualized two-dimensionally using a force-based algorithm (***Kamada and Kawai, 1989***) to determine node positions.

The small world index describes the efficiency of information flow in a network compared to a random network of the same size. Mathematically, the small world index (σ) is the quotient of a network’s average cluster coefficient normalized to the mean of 1000 random graphs with the same number of nodes and edged and its corresponding normalized average shortest path length. The higher the density of a network, the more it resembles a random network, and the smaller is σ. Therefore, σ was calculated over a wide range of network densities.

To define suitable density to compare network topology we determined the vertex of the fitted hyperbolic σ curve (formula 1) of the MS CTR group. The vertex of a hyperbola is the point where the curvature is maximum. For vertex determination, σ values were normalized to the range of densities to ensure the same data range on y and x axis. Mathematically, the maximum curvature can be defined as the point where the first derivative (formula 2) equates -1. Considering the resulting x-coordinate x_v_ (formula 3), the next highest full percent density was chosen for topological comparison.

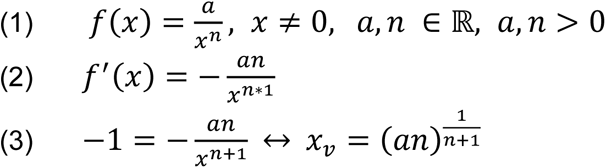

Additionally, to the characterization of the entire network, significant group differences in connectivity strength (i.e., edge weight) and several node-specific graph-theoretical parameters was calculated.

To detect significant differences in connectivity strength, Network-Based-Statistics (NBS) were used (***Zalesky et al., 2010***). Briefly, for two groups, a T-test was calculated on each connection, and the most significant component of contiguous connections with p < 0.05 were retained. To correct for multiple comparison, we performed permutation testing with building up the null distribution (across permutations) of the size (i.e., number of connections) of the most significant retained component. The estimated multiplicity-adjusted p-value (P_FWE_) for the original component was the proportion of permutation samples for which the component size was less than or equal to the particular original component size.

The nodes within the networks, which represent the activated brain regions, were characterized by following graph-theoretical parameters: *’strength,’ ‘degree,’ ‘clustering coefficient,’ ‘average path length,’ ‘betweenness centrality’* and *’hub score.’* Node *’strength’* is defined as sum of weights of all edges that are connected to that specific node, and *’degree’* as number of these edges. The *’clustering coefficient’* represents a measure of the degree to which nodes in a graph tend to cluster together. It is defined as the proportion of actual connections between all neighbors of a node, relative to all possible connections between these neighbors (***Bullmore and Sporns, 2009; Sporns, 2013b***). The dimension of *’average path length’* represents an indicator of efficiency of information flow in a network. For each node, they define it as average number of steps along the shortest paths of this node to all other network nodes (***Sporns et al., 2007***). Network hubs are nodes with high importance for the integrity of the whole network. Hubs usually are high degree nodes and connect distinct parts of a network. This property is characterized by ’*betweenness centrality,’* which quantifies the number of times a node acts as a bridge along the shortest path between two other nodes (***Freeman, 1977***). Hubs can be characterized by a *’hub score’*, a relative score assigned to all nodes in the network, based on the concept that connections to high-scoring nodes contribute more to the score of the node in question than equal connections to low-scoring nodes. Hyperlink-Induced Topic Search (HITS) (***Kleinberg, 1999***) was used to calculate the hub score. We normalized all parameters except for degree to 100 nodes within each network to adjust for network sizes. In networks with the same density, differences in degree showed differences in network size.

### Statistics

Each graph-theoretical parameter was statistically tested for significant effects using three-factor ANOVA with interactions separately for high (95 g / 5 mA) and low (50g / 1 mA) stimulation. The three factors were pain entity (INC, CFA, and SHA), stimulation modality (electrical and mechanical) and the activated brain regions. Main effects in the first two factors show a network (i.e., brain) wide modulation of the parameter, interaction with the factor brain regions a node (i.e. brain region) specific modulation. ANOVA main effects between groups were tested post-hoc using Tukey HSD with Bonferroni correction. Significant ANOVA interactions with factor brain regions were tested post-hoc with a homoscedastic t-test (FDR corrected with q=0.1) to identify group-specific differences per brain region.

In result, the complex high-dimensional set of graph-theoretical parameters characterized each node. In order to obtain dimensionality reduction and therefore a feature selection, a Linear Discriminant Analysis (LDA) was performed for high and low stimulation intensities, respectively, to classify nodes according to their group affiliation. LDA is a multivariate analysis that finds linear combinations of features which separates two or more groups. The resulting combinations represented a new coordinate system with reduced dimensionality and their dimensions ranked according to their power to separate groups. Therefore, the first three LDA dimensions were used to visualize group classification. Moreover, the feature loadings of the LDA dimension identified the features and their importance for the group separation, which corresponds to the respective biosignature.

In addition, for each node (i.e. brain region) we computed the Mahalanobis distance between its respective coordinates across all LDA dimensions in all group pairs. The higher the Mahalanobis distance, it separated the more this specific brain region between groups.

### Software

fMRI preprocessing and GLM analysis were performed with BrainVoyager OX 2.82 (Brain Innovation B.V. Maastricht, The Netherlands). For all other analysis steps including NBS, MagnAn 2.5 (BioCom, Uttenreuth), an IDL application (Exelis Visual Information Solutions Inc., a subsidiary of Harris Corporation, Melbourne, FL, USA) designed for complex image processing and analysis with emphasis on MR imaging, was used. ANOVA and t-tests were calculated with Real Statistics (http://www.real-statistics.com), a free statistic resource package for Excel 2013 (Microsoft® Office) and LDA in R (https://www.r-project.org), a free programming language for statistical calculations and graphics. Networks were visualized using AMIRA 5.4 (Thermo Fischer Scientific, Waltham, MA, USA) with custom modules.

