## Supplemental material for "Distinct Functional cerebral Hypersensitivity networks during incisional and inflammatory pain in rats"

Supplemental Figure 1

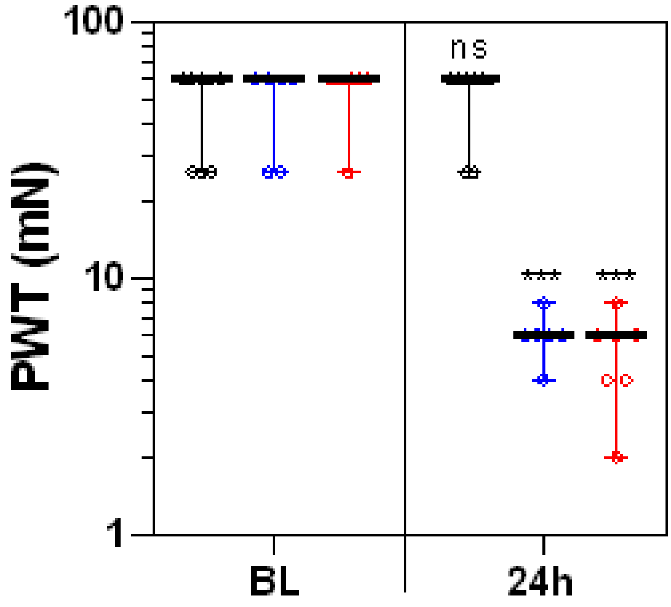

### Supplemental Figure 2

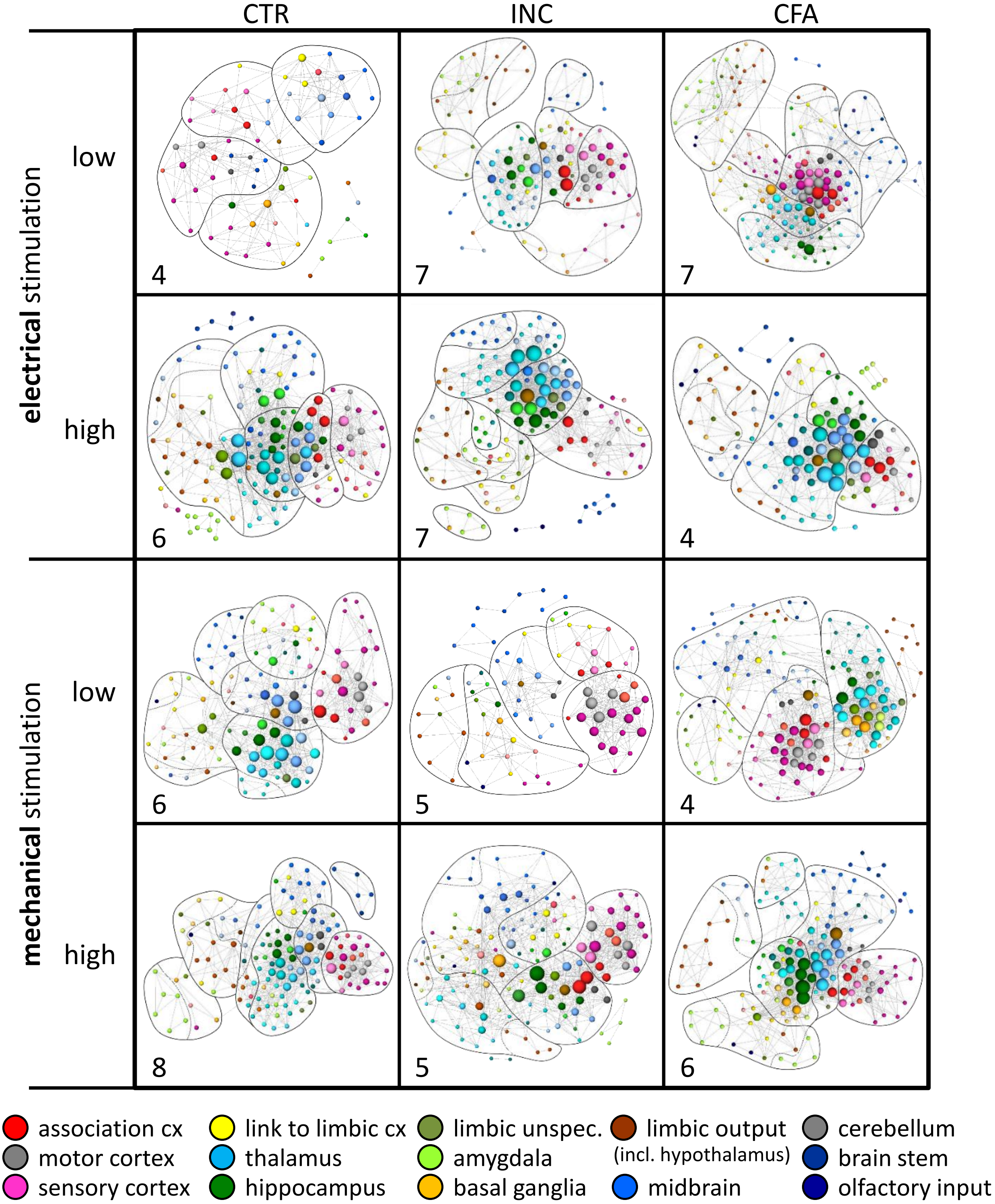

### Supplemental Figure 3

A

**low** stimulation intensity

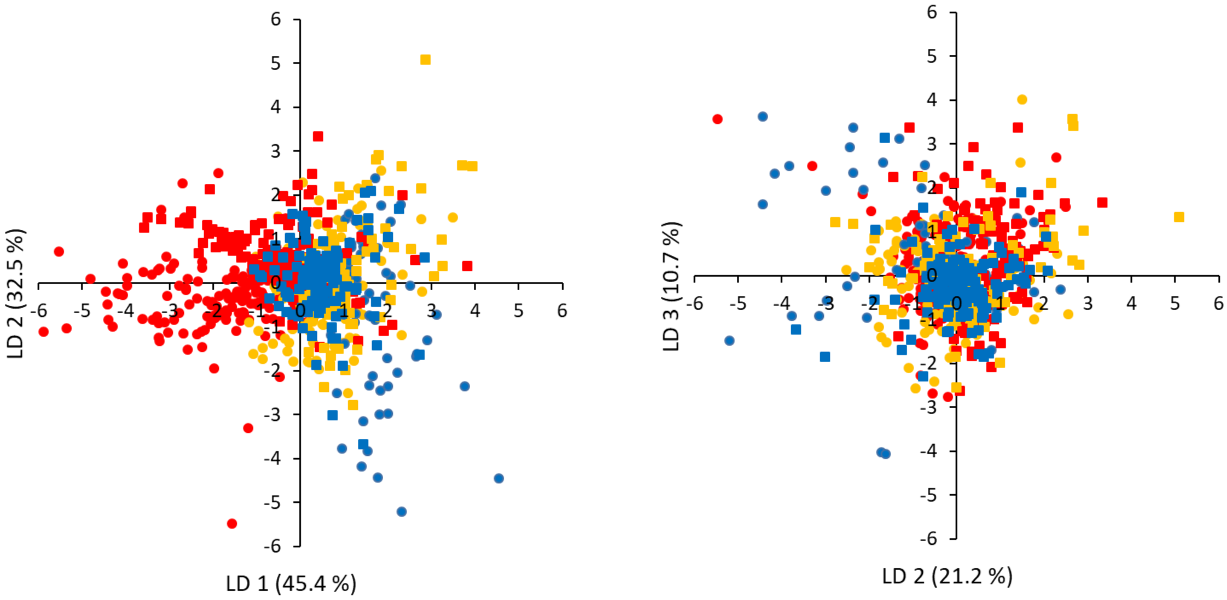


B

**high** stimulation intensity

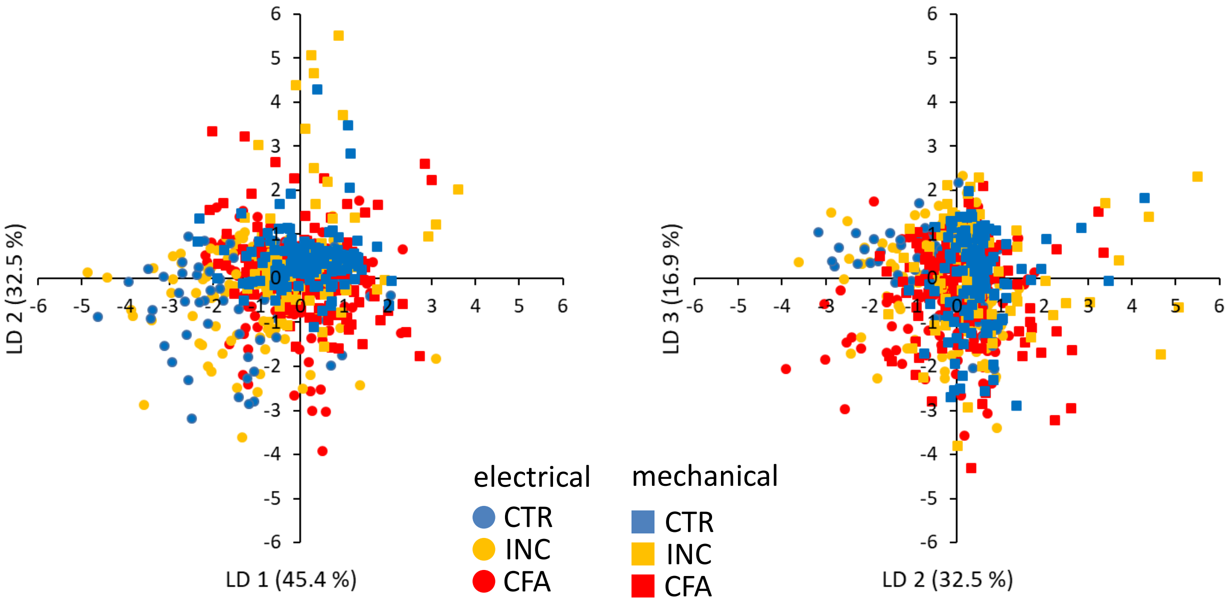

### Supplemental Figure 4

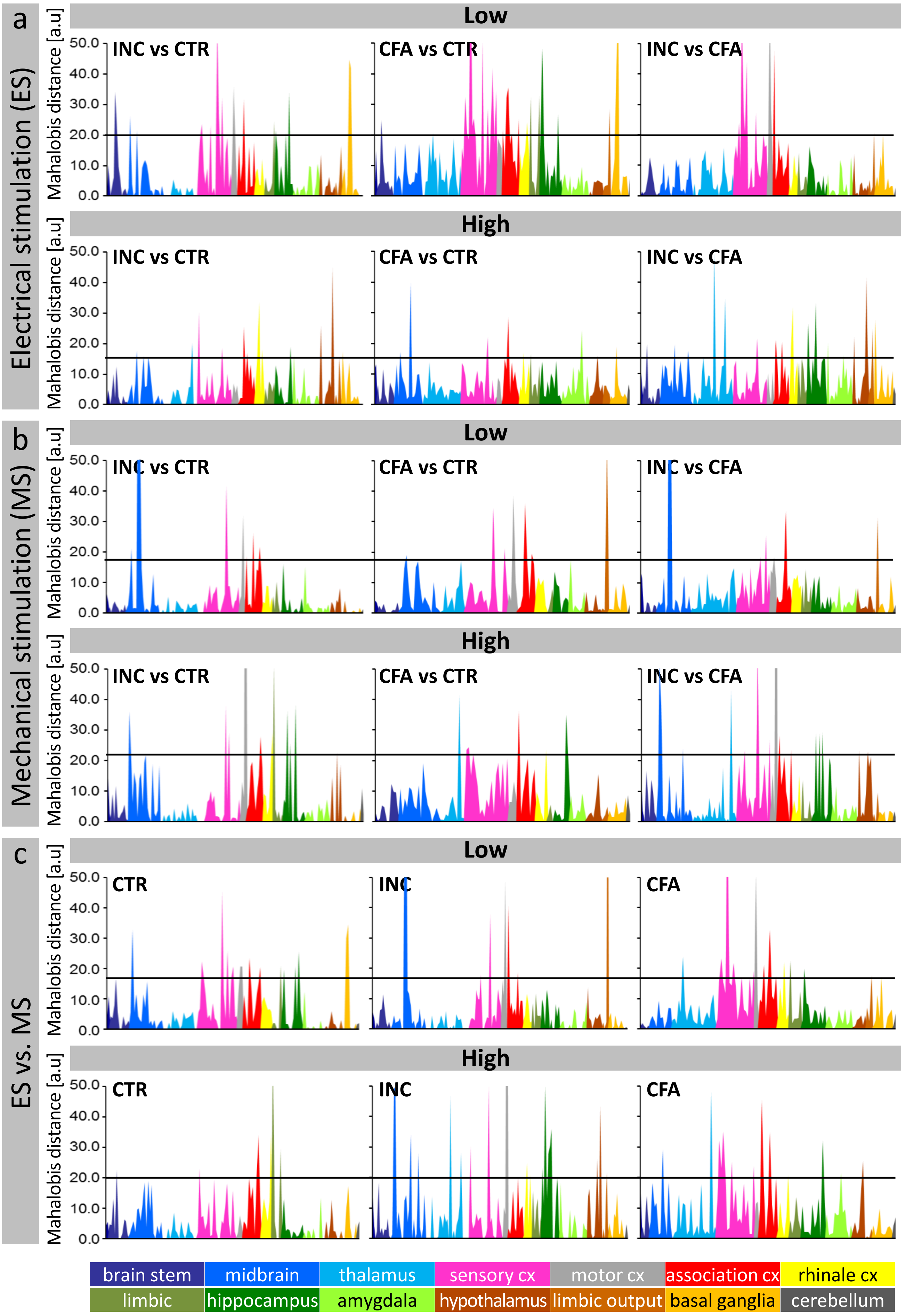

Supplemental Table 1: FDR threshold values (Pearson‘s correlation r) to define significant connections

| | | CTR | INC | CFA |
|---|---|---|---|---|
| **electrical stimulation** | **low** | 0.117 | 0.123 | 0.113 |
| | **high** | 0.104 | 0.104 | 0.111 |
| **mechanical stimulation** | **low** | 0.122 | 0.114 | 0.102 |
| | **high** | 0.126 | 0.117 | 0.102 |

Supplemental Table 2: Three Factor ANOVA

**low** stimulation intensity

| Factor | | Strength | Degree | Clustering coefficient | Average path length | Betweeness centrality | HIT hub score |
|---|---|---|---|---|---|---|---|
| Main effects | A Stimulation | **0.001** | 1.000 | 0.522 | 0.790 | 1.000 | 1.000 |
| | B Pain Entity | **<0.001** | **<0.001** | 1.000 | 1.000 | **<0.001** | 1.000 |
| | C Brain Region | **<0.001** | **<0.001** | **<0.001** | **<0.001** | **<0.001** | **<0.001** |
| Interactions | A x B | **0.012** | **<0.001** | 1.000 | **0.022** | **<0.001** | 1.000 |
| | A x C | 0.424 | 1.000 | **<0.001** | **<0.001** | 1.000 | **0.039** |
| | B x C | 0.330 | 1.000 | 0.340 | **<0.001** | 1.000 | 0.167 |
| | A x B x C | 1.000 | 1.000 | 0.211 | **<0.001** | 1.000 | **0.002** |

**high** stimulation intensity

| Factor | | Strength | Degree | Clustering coefficient | Average path length | Betweeness centrality | HIT hub score |
|---|---|---|---|---|---|---|---|
| Main effects | A Stimulation | 0.562 | **<0.001** | 0.940 | **<0.001** | **<0.001** | 1.000 |
| | B Pain Entity | **0.018** | **0.001** | 1.000 | 1.000 | **<0.001** | 1.000 |
| | C Brain Region | **<0.001** | **<0.001** | **<0.001** | **<0.001** | **<0.001** | **<0.001** |
| Interactions | A x B | **<0.001** | **<0.001** | 0.403 | **0.032** | **<0.001** | 1.000 |
| | A x C | **<0.001** | **<0.001** | **<0.001** | **<0.001** | **<0.001** | **<0.001** |
| | B x C | 1.000 | 1.000 | 0.100 | 1.000 | 1.000 | 1.000 |
| | A x B x C | **0.014** | 1.000 | 0.839 | 0.171 | 0.307 | 1.000 |

Bonferroni corrected p-values. Significant effects are highlighted in bold ($\alpha$=0.05)

Supplemental Table 4: List of brain regions including abbreviations and colors.

| Abbreviation | Brain region | Anatomical Group | Functional Group |
|---|---|---|---|
| ON | olfactory nuclei | olfactory input | olfactory input |
| OT | olfactory tubercle | olfactory input | olfactory input |
| AP | area postrema | medulla | brain stem |
| Sol | solitary tract | medulla | brain stem |
| MdD | dorsal medullary reticular nucleus | medulla | brain stem |
| MdV | ventral medullary reticular nucleus | medulla | brain stem |
| RtL | lateral reticular nucleus | reticular nucleus | brain stem |
| PCRt | parvicellular reticular nucleus | reticular nucleus | brain stem |
| Gi | gigantocellular reticular nucleus | gigantocellular nucleus | brain stem |
| PGiL | lateral paragigantocellular nucleus | gigantocellular nucleus | brain stem |
| R | raphe nucleus | raphe | brain stem |
| PnC | pontine reticular nucleus caudal | pons | brain stem |
| PnM | pontine reticular nucleus medial | pons | brain stem |
| PnO | pontine reticular nucleus oral | pons | brain stem |
| TN | tegmental nuclei | tegmentum | midbrain (sensory input) |
| VTA | ventral tegmental area | tegmentum | midbrain (sensory input) |
| PBL | lateral parabrachial nucleus | parabrachial nucleus | midbrain (sensory input) |
| CF | cuneiform nucleus | tegmentum | midbrain (sensory input) |
| Red | red nucleus | red nucleus | midbrain (sensory input) |
| IP | interpeduncular nucleus | interpendicular nucleus | midbrain (sensory input) |
| IC | inferior colliculus | colliculi | midbrain (sensory input) |
| SC | superior colliculus | colliculi | midbrain (sensory input) |
| PTA | pretectal area | pretectal Area | midbrain (sensory input) |
| SN | substantia nigra | mesencephalon | midbrain (sensory input) |
| Mes | mesencephalic region | mesencephalon | midbrain (sensory input) |
| thGM | medial geniculate nucleus | genigulate nucleus | thalamus |
| thGL | lateral geniculate nucleus | genigulate nucleus | thalamus |
| thGV | ventral geniculate nucleus | genigulate nucleus | thalamus |
| thLP | lateral posterior thalamic nucleus | lateral posterior thalamus | thalamus |
| thPo | posterior thalamic nuclear group | lateral thalamus | thalamus |
| thVM | ventromedial thalamic nucleus | lateral thalamus | thalamus |
| thVL | ventrolateral thalamic nucleus | lateral thalamus | thalamus |
| thVPM | ventral posteromedial thalamic nucleus | lateral thalamus | thalamus |
| thVPL | ventral postolateral thalamic nucleus | lateral thalamus | thalamus |
| thMD | mediodorsal thalamus | medial thalamus | thalamus |
| thLD | lateraldorsal thalamic nucleus | medial thalamus | thalamus |
| thSM | submedius thalamic nucleus | medial thalamus | thalamus |
| thA | anterior thalamic group | anterior thalamus | thalamus |
| thRt | reticular thalamic nucleus | reticular thalamus | thalamus |
| thRe | reuniens thalamic nucleus | reticular thalamus | thalamus |
| thPV | paraventricular thalamic nucleus | paraventricular thalamic nucleus | thalamus |
| cxS1HL | primary somatosensory cortex hind limb | somatosensory cortex | sensory cortex |
| cxS1FL | primary somatosensory cortex forelimb | somatosensory cortex | sensory cortex |
| cxS1DZ | primary somatosensory cortex dysgranular | somatosensory cortex | sensory cortex |
| cxS1J | primary somatosensory cortex jaw | somatosensory cortex | sensory cortex |
| cxS1ULp | primary somatosensory cortex upper lip | somatosensory cortex | sensory cortex |
| cxS1BF | primary somatosensory cortex barrel field | somatosensory cortex | sensory cortex |
| cxS1Tr | primary somatosensory cortex trunk | somatosensory cortex | sensory cortex |
| cxS1r | primary somatosensory cortex rest | somatosensory cortex | sensory cortex |
| cxS2 | secondary somatosensory cortex | somatosensory cortex | sensory cortex |
| cxAu | auditory cortex | auditory cortex | sensory cortex |
| cxVis | visual cotex | visual cortex | sensory cortex |
| cxPtA | parietal association cortex | parietal association cortex | association cortex |
| cxTeA | temporal association cortex | temporal association cortex | association cortex |
| cxRS | retrosplenial cortex | cingulate cortex | association cortex |
| cxCg | cingulate cortex | cingulate cortex | association cortex |
| cxPrL | prelimbic cortex | limbic association cortex | association cortex |
| cxIL | infralimbic cortex | limbic association cortex | association cortex |
| cxPdD | dorsal peduncular cortex | frontal association cortex | association cortex |
| cxOrb | orbital cortex | frontal association cortex | association cortex |
| cxFr3 | frontal cortex area 3 | frontal association cortex | association cortex |
| cxFrA | frontal association cortex | frontal association cortex | association cortex |
| cxIns | insular cortex | insular cortex | association cortex |

Supplemental Table 2 continued

| Abbreviation | Brain region | Anatomical Group | Functional Group |
|---|---|---|---|
| cxEnt | entorhinal cortex | rhinale cortex | link to the limbic |
| cxPrh/Ect | perirhinal/ectorhinal cortex | rhinale cortex | link to the limbic |
| cxPir | piriform cortex | piriform cortex | link to the limbic |
| Hb | habenuli | habenulae | limbic system unspecific |
| Sep | septum | septum | limbic system unspecific |
| DB | nuclei of diagonal band | diagonal band | limbic system unspecific |
| hcAD | anteriordorsaler hippocampus | dorsal hippocampus | hippocampus |
| hcPD | posteriordorsaler hippocampus | dorsal hippocampus | hippocampus |
| hcSD | dorsal subiculum | dorsal hippocampus | hippocampus |
| hcCA3v | ventral CA3 fields | Ca3 fields | hippocampus |
| hcV | ventraler hippocampus | ventral hippocampus | hippocampus |
| hcSV | ventral subiculum | ventral hippocampus | hippocampus |
| hcDGp | posterior layers of the dentate gyrus | dentate gyrus | hippocampus |
| amA | anterior amygdala | amygdala | amygdala |
| amM | medial amygdaloid nucleus | amygdala | amygdala |
| amCo | cortical amygdala | amygdala | amygdala |
| amBM | basomedial amygdaloid nucleus | amygdala | amygdala |
| amBL | basolateral amygdaloid nucleus | amygdala | amygdala |
| amCe | central nucleus of the amygdala | amygdala | amygdala |
| amHA | amygdala hip area | amygdala | amygdala |
| ampirTr | amygdala piriform transition rechts | amygdala | amygdala |
| amSLE | sublenticular extended amygdala | amygdala | amygdala |
| BNST | bed nucleus of stria terminalis | bed nucleus of stria terminalis | amygdala |
| hyM | medial hypothalamus | hypothalamus | limbic output |
| hyL | lateral hypothalamus | hypothalamus | limbic output |
| hyArc | arcuate hypothalamic nucleus | hypothalamus | limbic output |
| hyPV | paraventricular hypothalamic nucleus | hypothalamus | limbic output |
| hyDM | dorsomedial hypothalamus | hypothalamus | limbic output |
| hyPo | posterior hypothalamus | hypothalamus | limbic output |
| ZI | zona incerta | zona incerta | limbic output |
| PAG | periaqueductal gray | periaqueductal gray | limbic output |
| CoM | corpora mammillaria | corpora mammillaria | limbic output |
| CPu | caudate putamen | caudate putamen | basalganglia |
| AcbC | core subregion of the nucleus accumbens | nucleus accumbens | basalganglia |
| AcbSh | shell subregion of the nucleus accumbens | nucleus accumbens | basalganglia |
| GPL | lateral globus pallidus | Pallidum | basalganglia |
| VP | ventral pallidum | Pallidum | basalganglia |
| Cl | claustrum | claustrum | basalganglia |
| Cb | cerebellum | cerebellum | motor output |
| M2 | secondary motor cortex | motor cortex | motor output |
| M1 | primary motor cortex | motor cortex | motor output |
